# AGIcam: An Open-Source IoT-Based Camera System for Automated In-Field Phenotyping and Yield Prediction

**DOI:** 10.64898/2026.01.13.699185

**Authors:** Worasit Sangjan, Nisit Pukrongta, Trevor Buchanan, Arron H. Carter, Michael O. Pumphrey, Sindhuja Sankaran

## Abstract

Continuous, high-frequency monitoring is essential to capture rapid phenological transitions and dynamic crop responses to the environment. However, most phenotyping platforms lack the temporal resolution and automation required for consistent, season-long trait assessment. This study introduces AGIcam, an open-source IoT camera system for automated and continuous in-field plant phenotyping and yield prediction. The platform integrates solar-powered Raspberry Pi units with a modular software stack, comprising Node-RED, InfluxDB, Grafana, and Microsoft Azure, for automated data acquisition, transfer, and visualization. In the 2022 growing season, 18 AGIcam systems were deployed in spring and winter wheat breeding trials, maintaining an uptime of over 85% while capturing frequent RGB and NoIR imagery. Time-series vegetation indices derived from these images were used to predict yield using random forest and Long Short-Term Memory (LSTM) models. The LSTM approach achieved the highest accuracy approximately one week after heading, with mean prediction errors of 3.41% for spring wheat and 1.62% for winter wheat. These results highlight the potential of IoT-based platforms such as AGIcam to enable real-time, scalable, and effective phenotyping solutions for data-driven crop improvement.

## 1. Introduction

The Internet of Things (IoT) has become increasingly prominent in agriculture, offering automated, high-frequency monitoring capabilities essential for crop management and phenotyping. By linking distributed sensors with communication and cloud infrastructures, IoT platforms provide continuous in-field data streams that capture fine-scale crop responses to dynamically changing environments (Duguma & Bai, 2025). This persistent monitoring is crucial for breeding programs, where trait expression often depends on short-lived phenological transitions, and IoT systems enable traits to be tracked continuously throughout the season (Fan et al., 2021; Rajak et al., 2024; Hoyos-Villegas et al., 2025). Recent studies have demonstrated the potential of IoT-based phenotyping systems in maize and wheat, where cost-effective networks of imaging and environmental sensors improved data resolution and reduced the need for manual field visits (Numajiri et al., 2021; Fan et al., 2023; Pineda-Castro et al., 2024).

While IoT sensors provide plot-level data, other high-throughput phenotyping platforms, including unmanned aerial vehicles (UAVs), phenotyping towers, and satellites, offer broader spatial coverage (Marzougui et al., 2023; Shrestha et al., 2025). IoT-based systems, therefore, serve a complementary role by providing consistent temporal coverage and capturing local-scale crop responses that aerial or satellite observations may overlook. Their affordability and adaptability make them well-suited for multi-scale phenotyping frameworks, where integration of ground-based, aerial, and satellite data enables a more holistic understanding of genotype–environment interactions (Boursianis et al., 2022; Pinto et al., 2023; Liang et al., 2025).

Despite this promise, in-field deployment of IoT systems for crop phenotyping faces challenges that have slowed adoption in breeding programs. Remote trial locations often lack electricity and internet connectivity, making autonomous power systems and reliable communication networks essential for long-term operation (Sinha & Dhanalakshmi, 2022; Xu et al., 2022). Environmental stresses demand durable, low-maintenance hardware, while continuous imaging and sensor streams require efficient pipelines for storage, processing, and remote access. These barriers highlight the need for IoT-based systems that are autonomous, resilient, and capable of practical integration into breeding workflows (Farooq et al., 2020; Akhter & Sofi, 2022; Pathmudi et al., 2023).

One example is our earlier work, which introduced a Raspberry Pi–based camera system for wheat phenotyping (Sangjan et al., 2021). This system demonstrated the feasibility of using affordable, open-source hardware for continuous crop monitoring by delivering high-frequency image data suitable for deriving vegetation indices (VIs), which serve as key inputs for time-series trait prediction. While effective as a proof-of-concept, the system lacked power autonomy, connectivity, and automated data pipelines, which limited scalability. These constraints motivated the development of AGIcam (referring to the *Ag*riculture *I*oT *cam*era) system, a next-generation IoT platform capable of supporting season-long, real-time phenotyping under field conditions. Its architecture incorporates advances in IoT communication and wireless technologies that have been identified as critical for scaling agricultural monitoring and management (Tao et al., 2021; Tang et al., 2024).

Among the potential applications of IoT-derived VI data, yield prediction is one of the most widely studied, as VIs provide robust indicators of crop vigor, biomass, and final performance (Sangjan et al., 2023; Fan et al., 2024). Studies have shown that indices such as the Normalized Difference Vegetation Index (NDVI) and Green NDVI (GNDVI), when combined with machine learning and deep learning models such as random forest and Long Short-Term Memory (LSTM) networks, can achieve accurate yield estimation across crops and environments (Guo et al., 2024; Sangjan et al., 2024; Skobalski et al., 2024; Liu et al., 2025; Lu et al., 2025). However, most existing efforts have relied on single time points or aggregated indices, limiting their ability to capture temporal crop dynamics. IoT systems such as the AGIcam system address this limitation by generating continuous VI time series, which preserve fine-scale growth patterns and provide richer inputs for yield modeling. Evaluating random forest and LSTM models on these high-frequency datasets demonstrates the predictive potential of time-series approaches and the practical efficiency of IoT platforms in supporting yield prediction for breeding and crop management.

This study presents the design and field deployment of the AGIcam system, highlighting its hardware and software architecture, as well as its capability to generate high-frequency phenotypic data. Specifically, the study aims to: (1) describe the design and implementation of the AGIcam system as a scalable IoT platform for in-field phenotyping; (2) assess its ability to collect and transmit high-frequency phenotypic data across the season; and (3) evaluate the predictive performance of AGIcam-derived time-series data for yield estimation using machine and deep learning techniques. By integrating modular and open-source tools with real-time analytics, AGIcam offers a scalable and reproducible solution for high-resolution phenotyping, thereby advancing data-driven selection in modern breeding workflows.

## 2. Materials and Methods

### 2.1. Overview of Original System

The original camera platform was deployed during the 2020 wheat growing season as a proof-of-concept for automated field phenotyping (Sangjan et al., 2021). It was built around a Raspberry Pi Compute Module 3+ Lite connected to two Raspberry Pi Camera Module V2 units (8 MP), configured to capture synchronized Red-Green-Blue (RGB) and No-Infrared (NoIR) imagery. The dual-camera design enabled the calculation of VIs, such as NDVI and GNDVI, from paired visible and near-infrared channels, while RGB imagery supported traits like canopy cover and color dynamics.

Image acquisition was managed through custom Python scripts (https://www.python.org/, accessed May 9, 2025), with all data stored locally on microSD cards. A simple environmental sensor recorded air temperature and humidity, though this was not directly integrated into the imaging workflow. Data retrieval was manual, and subsequent VI calculation was performed offline to assess crop development. Further hardware specifications, calibration, and validation results are provided in Sangjan et al. (2021).

### 2.2. AGIcam System Hardware

AGIcam was developed as an enhanced version of the original dual-camera platform (Sangjan et al., 2021), with modifications for long-term, autonomous operation in field conditions. The following subsections describe the power supply system, wireless connectivity, and environmental sensor integration that distinguish this upgraded design.

#### 2.2.1. Power Supply System

Each AGIcam system was powered by a 6 W, 6 V Voltaic solar panel (Voltaic Systems, Brooklyn, NY, USA) connected to a 6,400 mAh Voltaic V25 USB battery through a 5 V controller that regulated charging and prevented overvoltage (Figure 1). The V25 battery was selected for its constant “always-on” output, which avoided shutdown under low loads and ensured operation of the Raspberry Pi and dual camera modules in the field.

**Figure 1.**
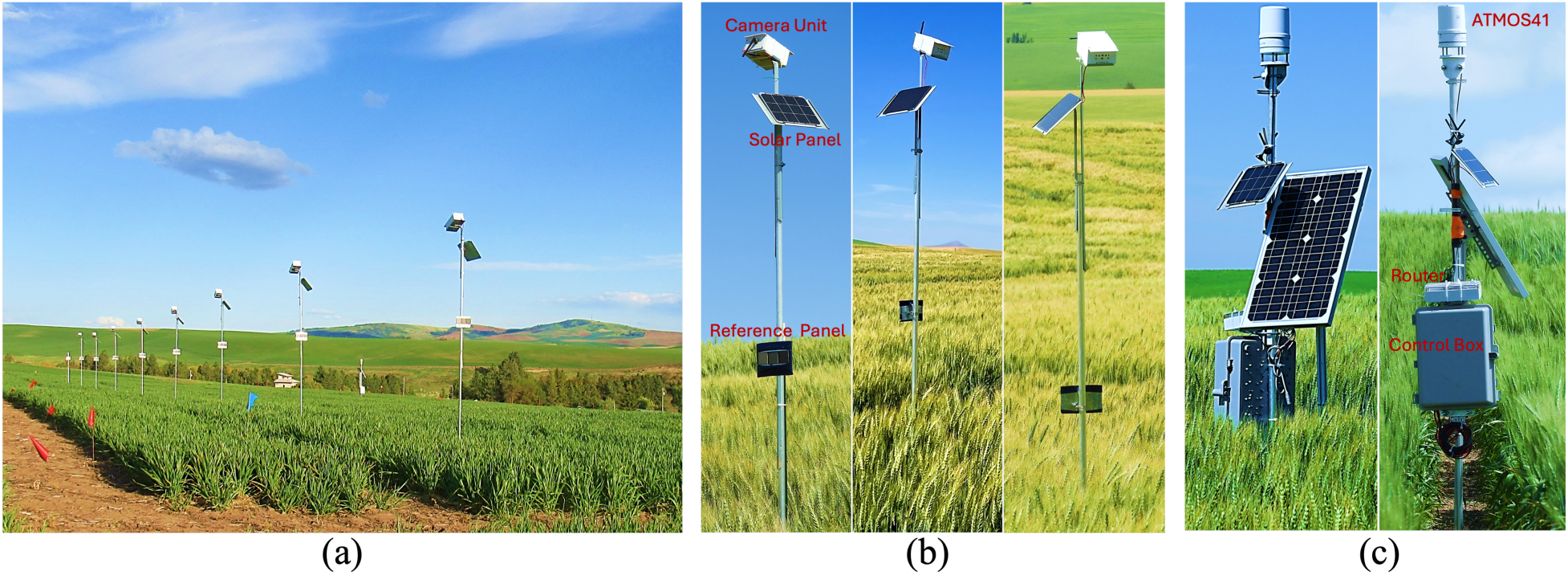
(a) AGIcam system deployed in a winter wheat breeding trial during the 2022 growing season; (b) individual AGIcam system; and (c) weather station and 4G LTE Wi-Fi router.

A Witty Pi power management board (UUGear, Prague, Czech Republic) was integrated to schedule power cycles of the Raspberry Pi, allowing the system to wake for image capture and then power down between sessions. This reduced unnecessary energy consumption and extended runtime during periods of low solar input. All components, including the Raspberry Pi, battery, and cameras, were enclosed in a weather-resistant, custom 3D-printed PLA (polylactic acid) housing for durability and easy field replacement (Figure 1b). The enclosure design files are openly available on GitHub (https://github.com/WorasitSangjan/IoT-based-Camera-Development, accessed May 9, 2025).

#### 2.2.2. Internet Connectivity

Each Raspberry Pi Compute Module 3+ Lite was equipped with a USB Wi-Fi antenna to enable wireless communication. Field-site internet access was provided by a 4G LTE Wi-Fi router (Linovision, Zhejiang, China), configured with a T-Mobile SIM card serving as the communication bridge for real-time data transfer and remote system access. The router was enclosed in a weatherproof housing and mounted on the control box (Figure 1c), which was powered by a 12 V DC battery recharged through a 30 W solar panel. The battery was placed inside the control box for environmental protection and stable operation (Figure 2).

**Figure 2.**
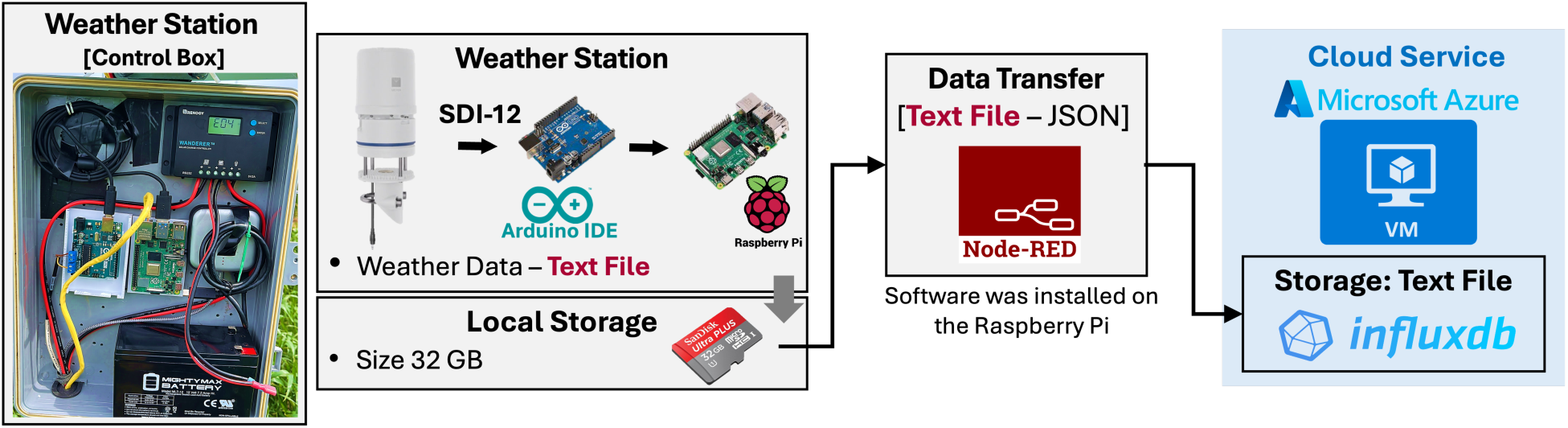
Schematic overview of the environmental monitoring system using the ATMOS41 weather station.

#### 2.2.3. Environmental Sensor Integration

Environmental data were recorded using an ATMOS41 all-in-one weather station (METER Group, Pullman, WA, USA) (Figure 1c), which measures 12 meteorological parameters, including air temperature, humidity, precipitation, wind speed, and solar radiation. The station communicated with an Arduino Uno (https://www.arduino.cc/, accessed May 9, 2025) through the SDI-12 protocol (1200 baud), a standard widely used for environmental sensors due to its low power demand and multi-drop capability. The Arduino relayed the data to a Raspberry Pi 4, which functioned as a data logger.

Data were transmitted over a USB serial connection, locally stored on an SD card, and uploaded using Node-RED (https://nodered.org/, accessed May 9, 2025) to a cloud-hosted InfluxDB database (https://www.influxdata.com/, accessed May 9, 2025) deployed on a Microsoft Azure virtual machine (https://azure.microsoft.com/en-us, accessed May 9, 2025) (Figure 2). This workflow mirrored the AGIcam system (Figure 3), enabling both environmental and image-derived time-series data to be integrated into a centralized database for management and analysis.

**Figure 3.**
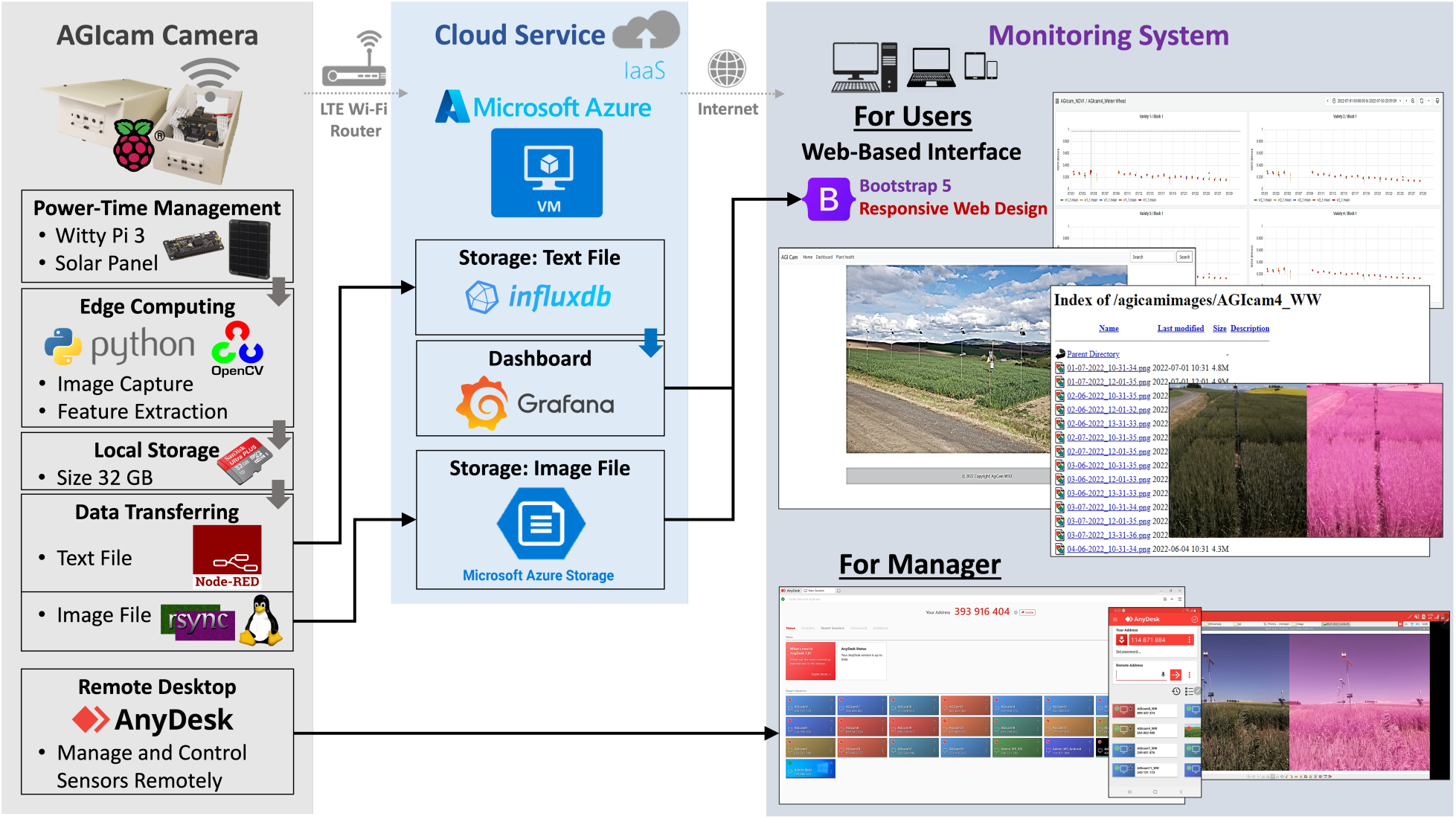
System architecture of the AGIcam system, illustrating its core components: field-deployed AGIcam system, cloud infrastructure, and user-facing monitoring interfaces. Abbreviations: IaaS - Infrastructure-as-a-Service; VM - Virtual Machine.

#### 2.2.4. Cost Summary

The total hardware cost per AGIcam system was approximately USD 150 to 200, depending on supplier availability. Core components included a Raspberry Pi board (∼ USD 35–50), dual RGB/NoIR camera modules (∼ USD 25–50), a Witty Pi 3 (∼ USD 10–15), SD card storage (∼ USD 5–10), and a solar panel with battery pack (∼ USD 60–80). A voltage regulator (∼ USD 5–10) and a custom 3D-printed enclosure (∼ USD 10–20) were also required. Mounting hardware (aluminum poles, base plates, and clamps) was added ∼ USD 30–40 per camera (Figure 1). Operational costs were modest, consisting of Microsoft Azure cloud services (∼ USD 3 per month) and a mobile internet service (∼ USD 30 per month per site). This cost structure enables the deployment of multiple cameras across distributed plots while maintaining long-term, continuous monitoring.

### 2.3. Backend System

The AGIcam’s backend operated on a modular, Linux-based software stack, which the system ran on the Raspbian Buster OS. All source code, including Python, JavaScript, and HTML scripts referenced in Sections 2.3, 2.4, and 2.6, is available on GitHub (https://github.com/WorasitSangjan/IoT-based-Camera-Development, accessed May 9, 2025).

#### 2.3.1. Camera Scheduling and Imaging Control

Power cycling of the Raspberry Pi was managed by the Witty Pi 3 Mini board, which activated the camera at a fixed time interval. Upon startup, a crontab executed Python scripts to capture synchronized RGB and NoIR images in PNG format (see Sangjan et al., 2021). An onboard Python pipeline processed each image to compute seven VIs (see Supplemental Material S1): Chlorophyll Index Green (CIgreen), Enhanced Vegetation Index 2 (EVI2), Green Normalized Difference Vegetation Index (GNDVI), Normalized Difference Vegetation Index (NDVI), Renormalized Difference Vegetation Index (RDVI), Soil Adjusted Vegetation Index (SAVI), and Simple Ratio (SR). These indices were selected based on the spectral capabilities of the dual RGB and NoIR camera system and their established relevance for yield prediction in cereal crops (Sandhu et al., 2021; Sangjan et al., 2024).

For each image and VI, summary statistics were calculated, including mean, median, maximum, standard deviation, and the 85th, 90th, and 95th percentiles. Percentiles were included to capture the canopy heterogeneity within the field of view, complementing average-based metrics. All processed outputs were saved with timestamp metadata in JSON format, written to the camera’s SD card, and queued for upload to the cloud.

#### 2.3.2. Data Transfer and Cloud Integration

Image and VI outputs were first stored locally on each AGIcam system’s SD card and then transmitted to the cloud using an Infrastructure-as-a-Service (IaaS) model, which provided scalable and secure storage of both unstructured (RGB/NoIR images) and structured (VI metrics) data (Figure 3). Raw images were transferred separately using the rsync (https://rsync.samba.org/, accessed May 9, 2025) over a Secure Shell (SSH) connection (Figure 3). Bandwidth and synchronization efficiency were maintained by configuring the system to transmit only one new image file per imaging session.

Time-series VI data were uploaded through a Node-RED workflow (Figure 4). At scheduled intervals, Node-RED was triggered after the imaging session and initiated data transfer by reading the locally generated vi.json file. The File Node loaded the VI dataset, which was then parsed into a JavaScript object via the JSON Node. A Function Node extracted VI statistics for each replicate–index pair, appended metadata such as plot ID and replicate ID, and formatted the payload into InfluxDB-compatible objects (measurement, fields, and tags). The structured payload was then routed to the corresponding InfluxDB bucket for cloud-based time-series storage. This modular workflow minimized transfer errors and ensured consistency of metadata across datasets.

**Figure 4.**
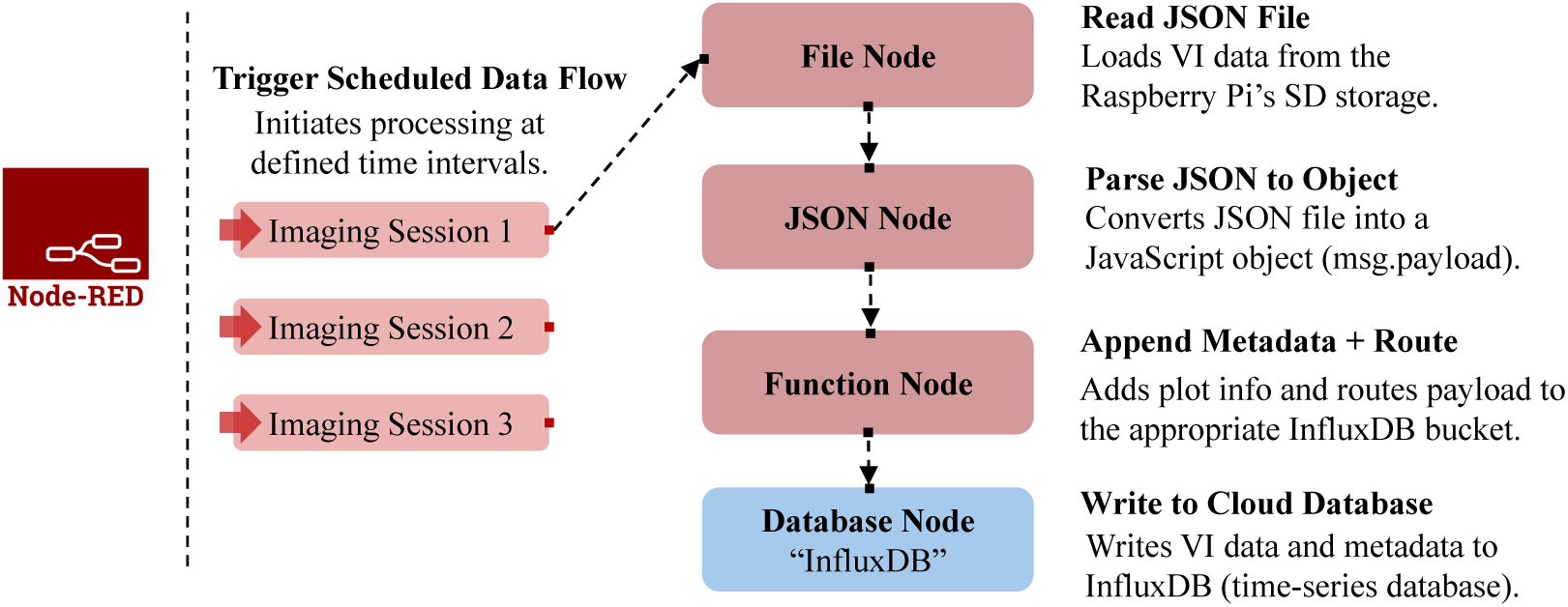
Node-RED data flow for automated transfer of vegetation index (VI) metrics to the InfluxDB cloud database.

### 2.4. Frontend System

A custom web interface was developed to support real-time visualization of AGIcam-derived data. The frontend consisted of modular HTML pages (index.html, dashboard.html) styled with Bootstrap 5 (https://getbootstrap.com/, accessed May 9, 2025) and hosted on the same Microsoft Azure virtual machine as the InfluxDB storage (Figure 3). This integration minimized latency between data logging and visualization, ensuring that processed results were accessible immediately.

Grafana (https://grafana.com/, accessed May 9, 2025) was used to generate interactive dashboards from time-series VI data stored in InfluxDB. These dashboards, embedded via iframes in the web interface and optimized for both desktop and mobile devices, enabled the dynamic visualization of NDVI and other indices over time (Figure 3). A separate page was allowed access to archived raw imagery collected by the AGIcam system.

Each AGIcam system was configured with AnyDesk (https://anydesk.com/en, accessed May 9, 2025) for remote desktop access (Figure 3). This setup enabled administrators or managers to perform software updates and troubleshooting without requiring on-site intervention, thereby reducing maintenance needs. Devices were linked to a managed account with fixed IDs and unattended access. A user manual describing dashboard navigation and remote-control setup is included in Supplemental Material S2.

### 2.5. Field Deployment and Data Collection

AGIcam systems were deployed during the 2022 growing season in spring and winter wheat (*Triticum aestivum* L.) breeding trials near Pullman, WA, USA. Trials followed a randomized complete block design. More details about the trial can be found in Sangjan et al. (2024). Twelve AGIcam systems were installed in spring wheat and six in winter wheat (Figure 5).

**Figure 5.**
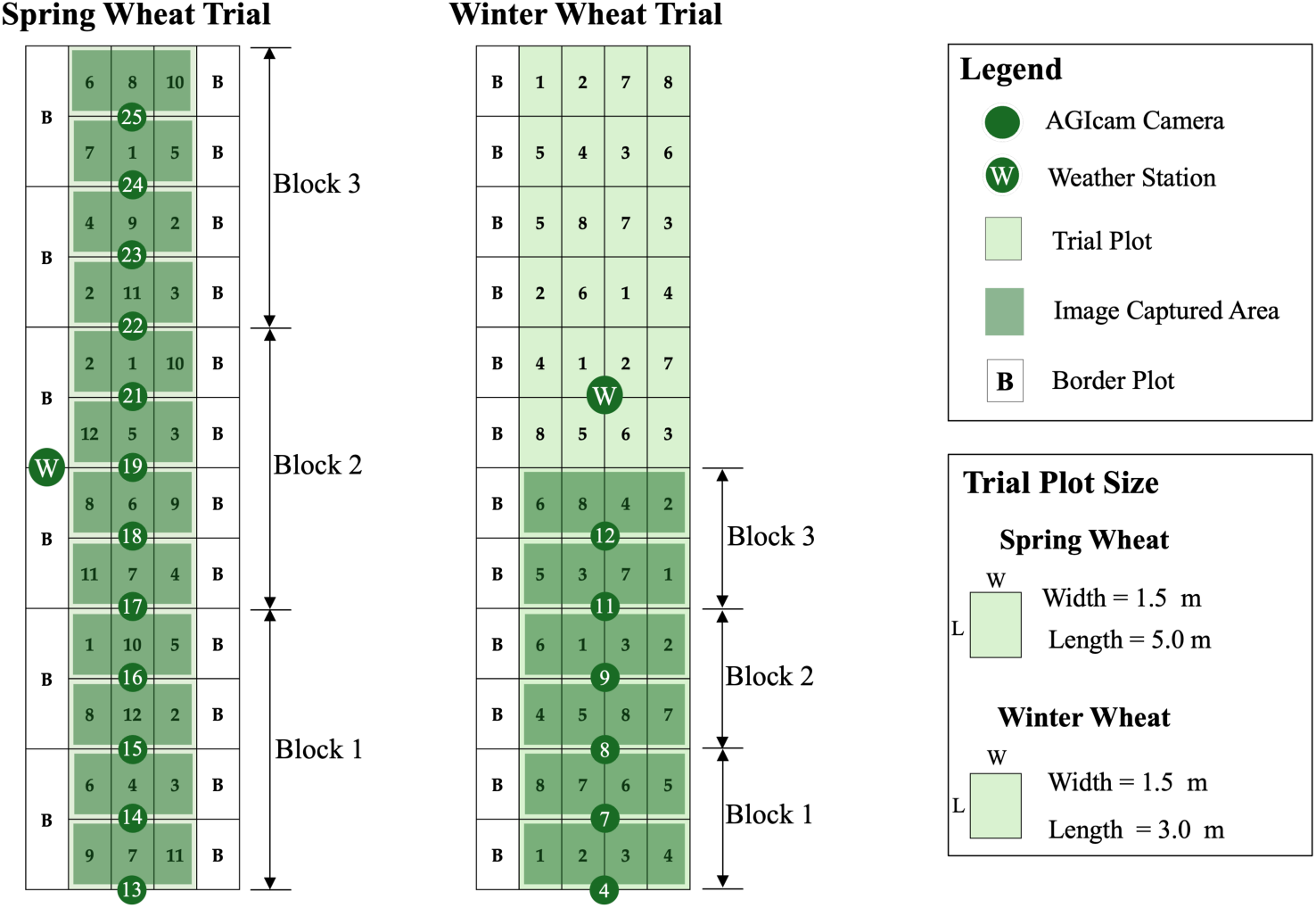
Plot layout and AGIcam system deployment for spring and winter wheat breeding trials during the 2022 season.

Each spring wheat AGIcam system monitored three adjacent plots (∼1.5 × 5.0 m, planted in eight rows), and each winter wheat AGIcam system monitored four adjacent plots (∼1.5 × 3.0 m, planted in seven rows). These dimensions reflect standard wheat breeding configurations in the Pacific Northwest, with three replicates of twelve varieties per trial (Figure 5). Cameras were positioned according to the field layout and oriented toward a reference panel to maintain consistent imaging geometry.

Each AGIcam system was mounted at a fixed height and angle to ensure full plot coverage and uniform image perspective. Installations followed the procedure outlined in Sangjan et al. (2021) and were co-located with an ATMOS41 weather station for environmental monitoring. Each camera captured synchronized RGB and NoIR images three times daily (at 10:30, 12:00, and 13:30), recording five images per session

### 2.6. Time-Series Phenotypic Data for Yield Prediction

Time-series VI data collected from the AGIcam system were used to predict grain yield in wheat breeding trials using random forest regression (Breiman, 2001) and LSTM networks (Hochreiter & Schmidhuber, 1997). Random forest was utilized for identifying critical growth windows because of its ability to model nonlinear relationships and assess feature importance (Figure 6). LSTMs were applied to capture sequential patterns and temporal dependencies in crop development (Silva et al., 2023; Tian et al., 2021). This analysis evaluated the predictive value of AGIcam-derived time series and demonstrated the system’s utility for downstream phenotyping applications.

**Figure 6.**
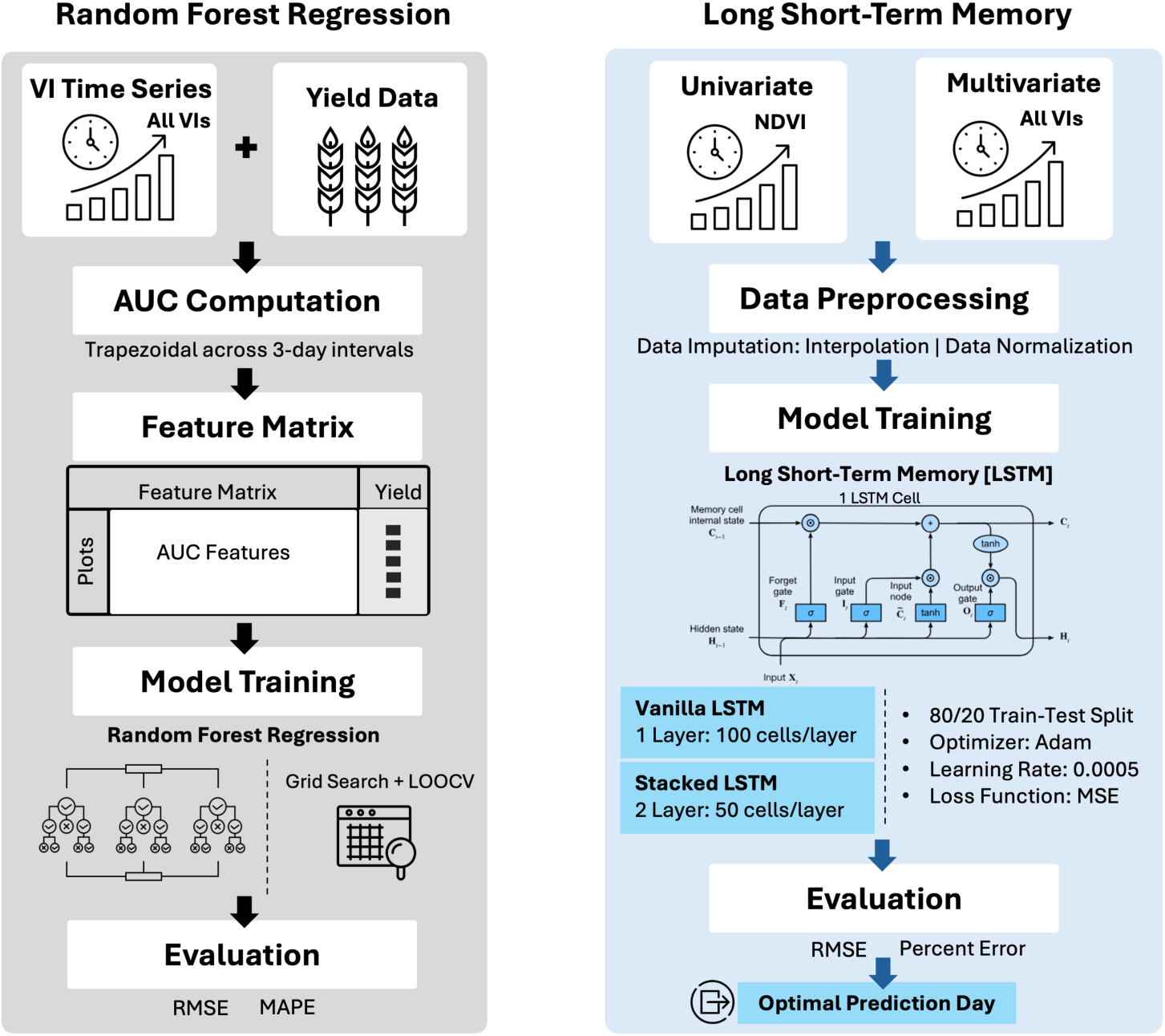
Machine learning and deep learning frameworks for yield prediction using time-series vegetation index (VI) data. Abbreviations: NDVI - Normalized Difference Vegetation Index; AUC - Area Under the Curve; LOOCV - Leave-One-Out Cross-Validation; RMSE - Root Mean Square Error; MAE - Mean Absolute Error; MSE - Mean Squared Error.

#### 2.6.1. Random Forest Regression

Seven VIs were aggregated into area under the curve (AUC) features using the trapezoidal rule (Iserles, 1996). The time series was divided into an initial seven-day window for early-season variability, followed by non-overlapping three-day intervals (e.g., Days 8–10, 11–13), yielding 19 intervals for spring wheat and 25 intervals for winter wheat. This structure preserves crop growth dynamics while reducing dimensionality, facilitating random forest modeling (Figure 6).

AUC features were paired with plot-level yield data and modeled using the RandomForestRegressor module from scikit-learn (https://scikit-learn.org/stable/, accessed May 9, 2025). Hyperparameters, including the number of trees, maximum tree depth, and the minimum split threshold, were optimized via a grid search with leave-one-out cross-validation (LOOCV) across ten randomized loops. Model performance was evaluated by using the root mean square error (RMSE) and the mean absolute percentage error (MAPE). Feature importance scores, averaged across folds, were used to identify the most predictive VIs and growth states for yield estimation.

#### 2.6.2. Long Short-Term Memory

LSTM models were implemented to model temporal dependencies in VI time series (Figure 6). Two input configurations were tested: univariate (NDVI only) and multivariate (all seven VIs). Missing values were linearly interpolated, and sequences were normalized before training. Padded timesteps introduced during sequence alignment were masked during model training, ensuring they did not bias predictions.

Two architectures were tested in TensorFlow (https://www.tensorflow.org/, accessed May 9, 2025). The first model was a Vanilla LSTM with a single 100-unit layer, using ReLU activation and L2 regularization (λ = 0.01). The second was a Stacked LSTM with two LSTM layers of 50 units (ReLU activation, L2 = 0.01), where the first layer returned sequences to the second. Both models incorporated batch normalization, dropout (0.2), and a dense output layer. Training used the Adam optimizer (learning rate = 0.0005) with mean squared error (MSE) as the loss function, an 80/20 train-test split, and early stopping (patience = 75) based on validation loss.

Model performance was evaluated using RMSE and percent error across the prediction timeline. For each plot, the optimal prediction day was defined as the day with the lowest RMSE, and overall performance was summarized as the mean RMSE and percentage error across all plots.

## 3. Results and Discussion

### 3.1. AGIcam System Performance and Reliability

During the 2022 growing season, AGIcam systems operated autonomously in both spring and winter wheat trials, capturing images three times per day as scheduled. The solar-powered configuration supported continuous monitoring for approximately two months in spring wheat and three months in winter wheat, consistent with the field deployment described in Section 2.5.

Most AGIcam systems retained uptime above 85%, with mean values of 89.9% for spring wheat and 88.8% for winter wheat (Figure 7). A temporary power outage early in the spring season, caused by several consecutive cloudy days, highlighted the importance of energy resilience in less sun-abundant environments. Although quickly resolved through remote monitoring via AnyDesk, this event suggests that larger batteries, additional solar panel capacity, or adaptive power-saving routines may improve future deployments.

**Figure 7.**
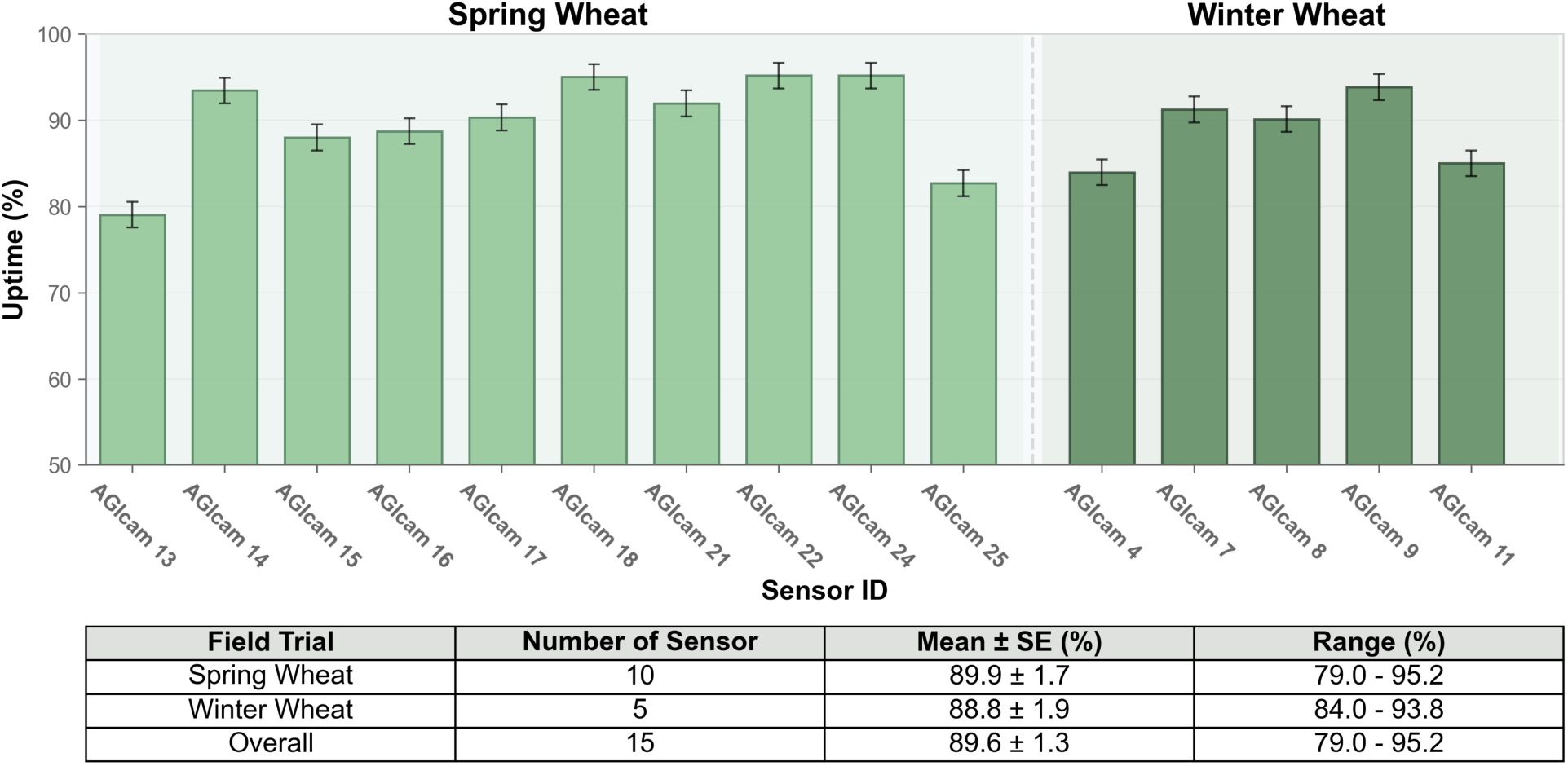
Uptime performance of AGIcam systems deployed in spring and winter wheat field trials during the 2022 growing season.

Aside from this isolated incident, the AGIcam system functioned reliably. The protective enclosure preserved system integrity, and the local execution of image capture, VI extraction, and metadata generation minimized backend dependencies. Integration with the cloud infrastructure enabled low maintenance and near real-time operation. Notably, the system also replaced commercial weather station loggers with an open-source alternative using the same backend. Three cameras (AGIcam 12, 19, and 23) experienced hardware failures, resulting in prolonged data loss, which underscores the value of redundancy planning, such as having spare units and a modular design, to ensure continuity during extended trials.

### 3.2. Wireless Connectivity and Network Limitations

Although the 4G LTE router setup described in Section 2.2.2 generally provided stable connectivity across most AGIcam systems and weather stations, performance declined for cameras located more than 50 m from the router. This resulted in occasional synchronization delays and missed uploads, particularly during periods with reduced solar power. These issues contributed to the lower uptime observed for a few cameras (Figure 7) and highlighted the sensitivity of wireless performance to both signal strength and environmental conditions. In particular, limited cellular band coverage at the site emphasized the importance of evaluating service providers during deployment planning, especially in remote agricultural sites.

Despite these challenges, the majority of cameras maintained consistent data transfer throughout the season. Future deployments could benefit from improved networking infrastructure, such as high-gain directional antennas, mesh Wi-Fi extenders, or long-range protocols like LoRa WAN (Long Range Wide Area Network) (Wang et al., 2024). Where available, 5G networks may offer increased bandwidth and broader coverage to support scalable, multi-AGIcam system deployments across larger or more remote trial locations (Kasera et al., 2024). In regions lacking terrestrial network access, satellite internet, particularly via low-Earth orbit systems, offers a viable alternative for real-time connectivity, albeit at a higher cost and with greater energy requirements (Ramírez-Arroyo et al., 2025).

### 3.3. Backend Automation and Data Flow Stability

The AGIcam system backend pipeline, comprising automated image capture, VI extraction, JSON generation, and cloud upload, operated with minimal manual intervention throughout the season. Python-based routines executed reliably, and the modular design with Node-RED and rsync supported low-latency data transfer (Figure 4).

Configuring Node-RED flows required manual linking of VI outputs to plot metadata, a step that was time-consuming and prone to labeling errors. Future iterations could integrate GPS (Global Positioning System)-based plot coordinates or scannable identifiers (e.g., QR codes) during imaging to streamline metadata association and improve traceability. Although only one image per session was retained, seasonal data volume still reached several gigabytes per camera. Cloud storage on Microsoft Azure was sufficient; however, further efficiency could be achieved through edge-based filtering, selective retention, or compression, which would reduce storage and bandwidth demands without compromising analytical value.

Data security was supported through open-source protocols. Node-RED parsed and routed VI data into InfluxDB, while rsync over SSH ensured encrypted image synchronization. Future deployments could incorporate firewall rules and user authentication to safeguard camera endpoints, providing protection comparable to commercial platforms while maintaining an open-source architecture.

### 3.4. Frontend Visualization and Usability

The frontend system provided reliable, near real-time monitoring of phenotypic trends throughout the 2022 growing season (Figure 3). Interactive Grafana dashboards displayed NDVI time-series data from all operational AGIcam systems, and users accessed raw image archives through the web interface (Figure 8). The system functioned smoothly on both desktop and mobile devices, with no major performance issues reported during deployment.

**Figure 8.**
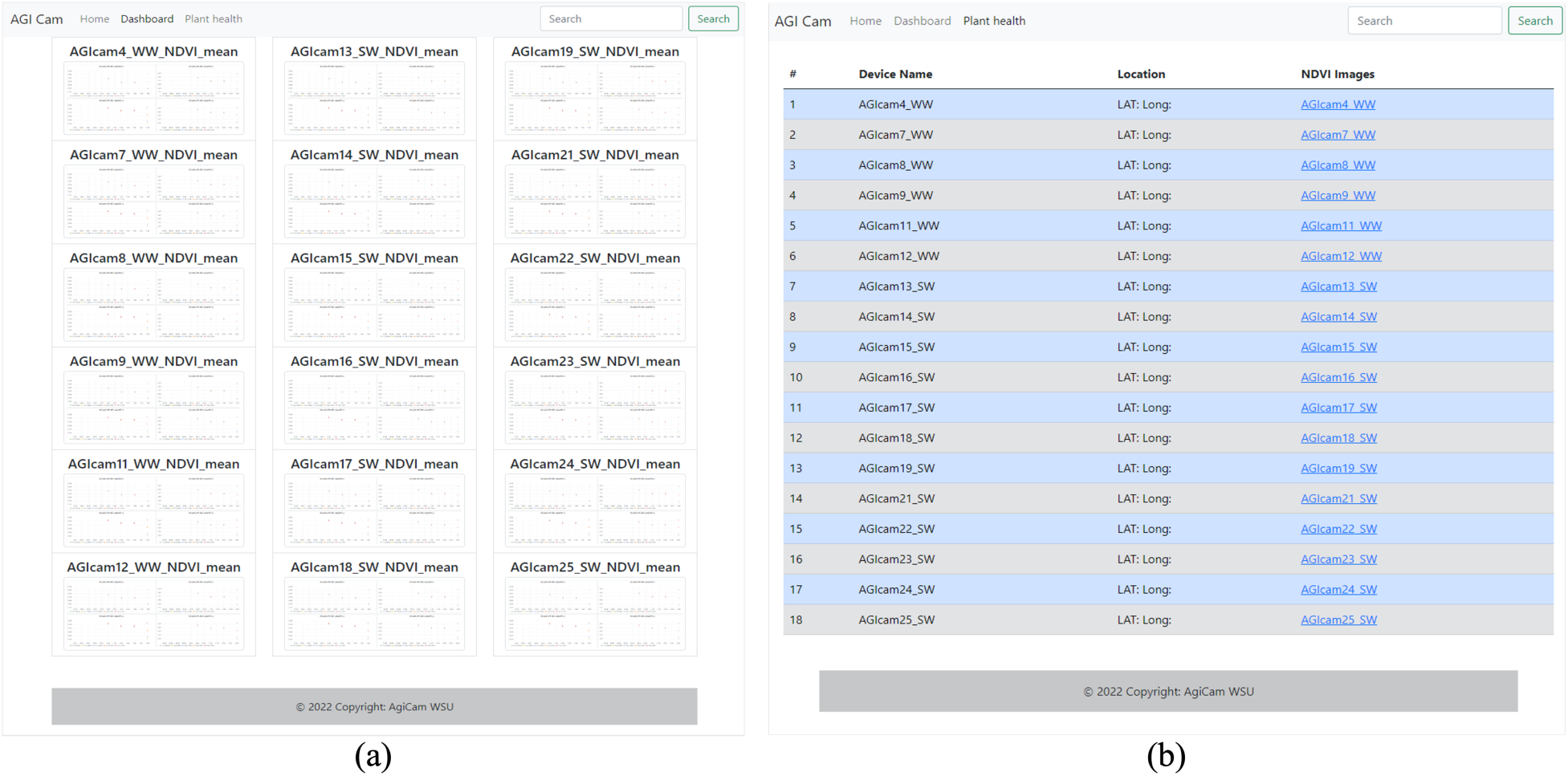
A web-based AGIcam dashboard interface was used for real-time visualization and monitoring during the 2022 growing season. (a) Dashboard view displaying time-series NDVI plots for each camera, organized by device ID and trial. (b) “Plant Health” page listing all AGIcam systems and hyperlinks to visual and downloadable imagery. Abbreviations: NDVI - Normalized Difference Vegetation Index; SW - Spring Wheat; WW - Winter Wheat.

Remote management via AnyDesk (Figure 3) enabled rapid troubleshooting, reducing downtime in the event of rare software or connectivity interruptions. While the core functionality was practical, several improvements could enhance usability for breeding applications, including expanding support to additional VIs, adding customizable metric selection, and implementing user-level access controls. The initial AnyDesk configuration, which relied on fixed IDs without session passwords, also introduced minor security risks; adopting PIN-based authentication and encrypted sessions would strengthen cybersecurity in future versions.

### 3.5. Yield Prediction Using Time-Series Vegetation Index Data

This section evaluates the utility of AGIcam-derived time-series VI data for yield prediction using machine learning and deep learning models.

#### 3.5.1. Random Forest Regression

Random forest models based on AUC features of VI time series (Figures 9a–b) achieved their highest accuracy around the heading stage in both spring and winter wheat trials (Figures 9c–d). For spring wheat, prediction error was lowest two days before heading (RMSE = 514 kg/ha; MAPE = 8.11%), whereas winter wheat reached its minimum error nine days before heading (RMSE = 984 kg/ha; MAPE = 9.31%).

**Figure 9.**
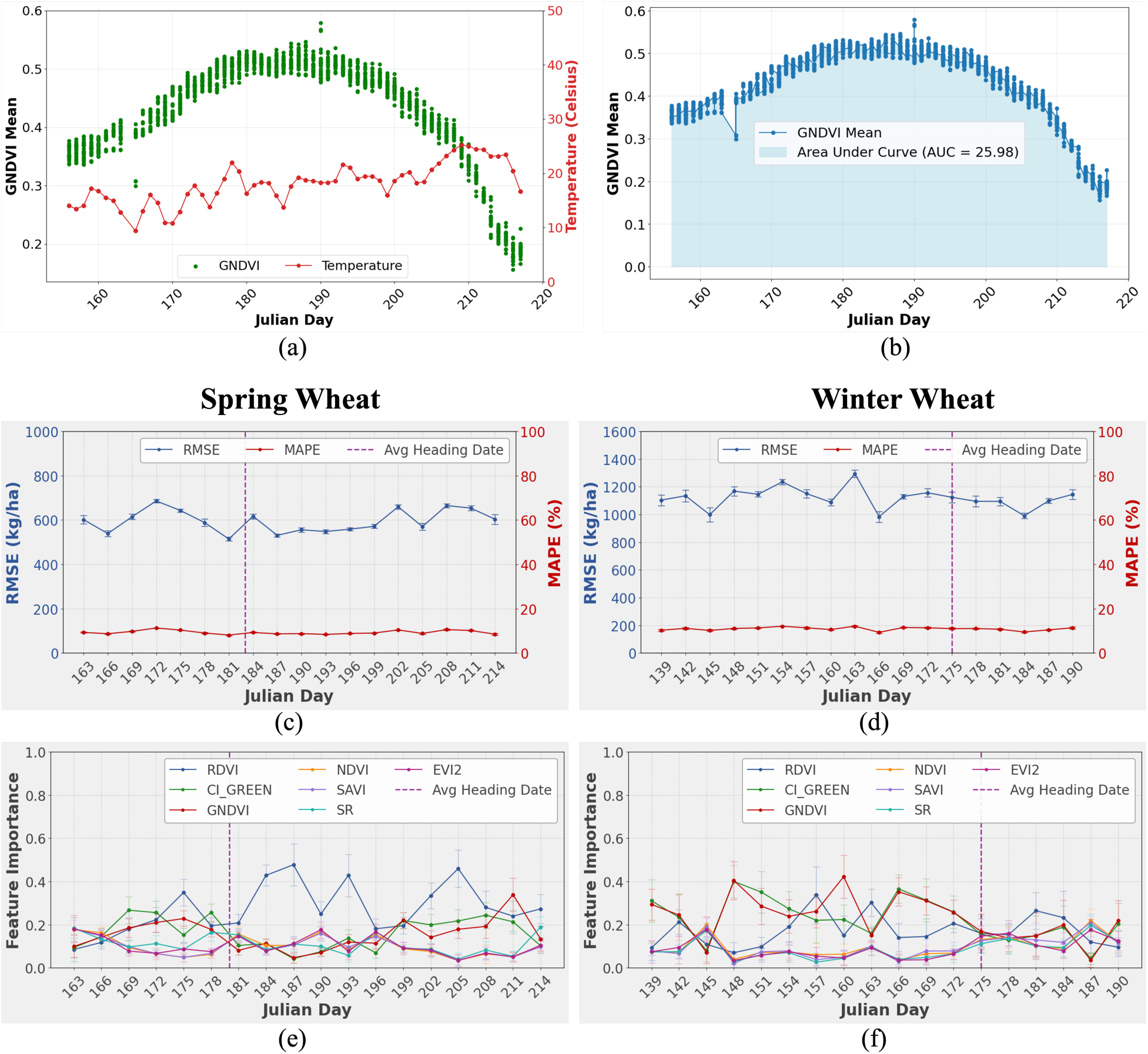
Yield prediction inputs and random forest model performance using time-series vegetation index (VI) data. (a) Daily GNDVI and corresponding air temperature profile of the 2022 growing season for a representative spring wheat trial. (b) Area under the curve (AUC) of the GNDVI profile from the same trial. (c, d) Prediction error trends for spring and winter wheat trials, showing root mean square error (RMSE) and mean absolute percentage error (MAPE) across Julian days. (e, f) Feature importance across the growing season for seven VIs in spring and winter wheat trials. Abbreviations: CIgreen - Chlorophyll Index Green; EVI2 – Enhanced Vegetation Index 2; GNDVI - Green Normalized Difference Vegetation Index; NDVI - Normalized Difference Vegetation Index; RDVI - Renormalized Difference Vegetation Index; SAVI - Soil Adjusted Vegetation Index; SR - Simple Ratio.

Model accuracy remained stable during the first week after heading, with RMSE values averaging 545 kg/ha (MAPE = 8.60%) in spring wheat and 1,060 kg/ha (MAPE = 10.41%) in winter wheat. This stability highlighted the biological importance of the heading-to-early grain filling window as a reliable stage for yield prediction using VI data.

Feature importance analysis identified CIgreen, GNDVI, and RDVI as the most influential indices during the heading (Figures 9e–f). These indices capture canopy chlorophyll content and structural dynamics, consistent with their frequent role as strong predictors of yield in prior UAV- and satellite-based studies (e.g., Liu et al., 2022; Segarra et al., 2022; Sangjan et al., 2024).

#### 3.5.2. Long Short-Term Memory

LSTM models were trained using AGIcam-derived VI time-series under both univariate (NDVI only) and multivariate (seven VIs) configurations. For spring wheat, the optimal setup was a Vanilla LSTM with NDVI as the sole input, whereas winter wheat achieved best performance with a multivariate LSTM (Figure 10).

**Figure 10.**
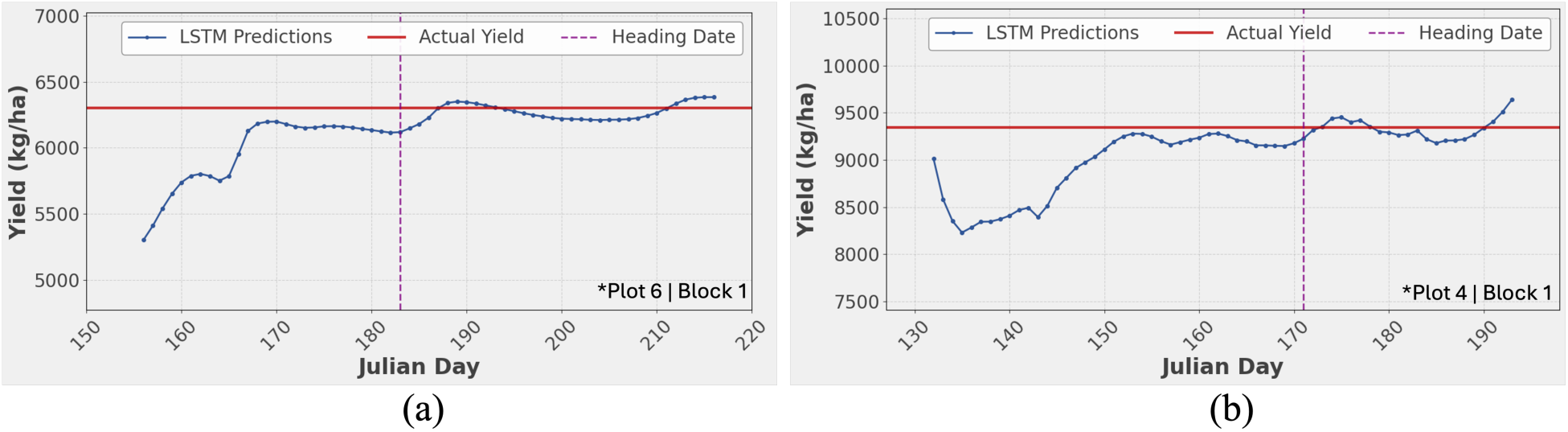
Daily yield predictions from Long Short-Term Memory (LSTM) models for individual plots of (a) spring wheat and (b) winter wheat. *Plot locations correspond to those shown in Figure 5.

Evaluation on a per-plot basis, using the day with the lowest RMSE as the optimal prediction date, accounted for phenological variation across genotypes. Most plots reached peak accuracy within a week after heading, coinciding with the early grain filling stage. At these time periods, spring wheat achieved an average RMSE of 222 kg/ha (3.41% error), and winter wheat 210 kg/ha (1.62% error). Representative predictions are shown in Figure 10.

These results mirrored the temporal patterns observed with random forest (Section 3.5.1), with both methods identifying the post-heading period as most predictive. However, the LSTM outperformed the random forest overall, reflecting its capacity to capture sequential dependencies in high-frequency time series. Prior satellite-based studies reported higher RMSEs (240–340 kg/ha; Cao et al., 2021; Cheng et al., 2022), likely due to lower temporal resolution. The sub-daily VI data from AGIcam systems provided finer phenological coverage, demonstrating the potential of IoT-based imaging for accurate, in-season yield forecasting.

### 3.6. Broader Applications and Future Perspectives

Beyond yield prediction from VI time series, the RGB and NoIR imagery collected by the AGIcam system provides opportunities for advanced image-based phenotyping. The fixed imaging geometry and consistent field of view across time make these datasets suitable for object detection tasks, including plant counting, wheat head detecting, and early disease symptom identification (Ahmad et al., 2023; Sanaeifar et al., 2023; Sangjan et al., 2025a). Such applications could expand spatial phenotyping and enable plot-level trait quantification at both fine spatial and temporal scales.

AGIcam data can also be integrated with environmental variables to support hybrid modeling approaches. Coupling image-derived traits with weather dynamics may improve prediction robustness under variable environments (Sangjan et al., 2025b). This highlights AGIcam’s role not only as a VI monitoring system, but also as a flexible data platform for broader phenomics analysis and breeding decision support.

Although the AGIcam system demonstrated effectiveness under standard field conditions, broader deployments may face challenges such as limited solar availability or reduced connectivity in remote sites. Addressing these operational constraints and validating the system across additional crops and seasons will be crucial for scaling up adoption and ensuring its utility in diverse agricultural contexts.

## 4. Conclusion

The AGIcam system was developed as an affordable, open-source IoT platform for automated in-field phenotyping. The integration of solar-powered hardware, modular software, and cloud connectivity enabled reliable, high-frequency monitoring tailored to breeding trials. Field deployment in wheat trials demonstrated stable operation and consistent data acquisition. Time-series VI data generated by the AGIcam system supported accurate yield prediction, with LSTM models outperforming random forest by capturing temporal crop dynamics. These findings demonstrate AGIcam’s utility as a scalable tool for yield forecasting and in-season monitoring. Nevertheless, the study was limited to a single growing season and two classes of wheat, underscoring the need for broader testing across crops, environments, and seasons. Future efforts should expand toward real-time trait detection and multimodal data fusion, positioning AGIcam as a flexible platform for phenomics and breeding decision support.

## CRediT authorship contribution statement

**Worasit Sangjan**: Conceptualization, Data curation, Formal analysis, Funding acquisition, Investigation, Methodology, Project administration, Resources, Software, Validation, Visualization, Writing – original draft, Writing – review & editing. **Nisit Pukrongta**: Methodology, Software, Writing – review & editing. **Trevor Buchanan**: Formal Analysis, Software, Writing – review & editing, **Arron H. Carter**: Funding acquisition, Investigation, Supervision, Resources, Validation, Writing – review & editing. **Michael O. Pumphrey**: Funding acquisition, Investigation, Supervision, Resources, Validation, Writing – review & editing. **Sindhuja Sankaran**: Conceptualization, Funding acquisition, Supervision, Investigation, Project administration, Resources, Validation, Writing – review & editing.

## Data availability

The source code and documentation for AGIcam are available on GitHub (https://github.com/WorasitSangjan/IoT-based-Camera-Development, accessed on May 9, 2025). The data is available at: https://doi.org/10.5281/zenodo.17970104.

## Acknowledgments

This study was funded by the United States Department of Agriculture (USDA) - National Institute for Food and Agriculture (NIFA) competitive project (accession number 1028108), AgAID Institute (2021-67021-35344), hatch project (accession number 1014919), and Washington State University’s College of Agricultural, Human, and Natural Resource Sciences’ Emerging Research Issues competitive grant opportunity (ERI-20-04). The authors would like to thank Dr. Milton Valencia Ortiz and Kingsley Charles Umani for their support during the field camera installation.

## References

Ahmad, A., Saraswat, D., & El Gamal, A. (2023). A survey on using deep learning techniques for plant disease diagnosis and recommendations for development of appropriate tools. Smart Agricultural Technology, 3, 100083. 10.1016/j.atech.2022.100083

Akhter, R., & Sofi, S. A. (2022). Precision agriculture using IoT data analytics and machine learning. Journal of King Saud University - Computer and Information Sciences, 34(8), 5602–5618. 10.1016/j.jksuci.2021.05.013

Boursianis, A. D., Papadopoulou, M. S., Diamantoulakis, P., Liopa-Tsakalidi, A., Barouchas, P., Salahas, G., Karagiannidis, G., Wan, S., & Goudos, S. K. (2022). Internet of Things (IoT) and Agricultural Unmanned Aerial Vehicles (UAVs) in smart farming: A comprehensive review. Internet of Things, 18, 100187. 10.1016/j.iot.2020.100187

Breiman, L. (2001). Random forests. Machine Learning, 45(1), 5–32. 10.1023/A:1010933404324

Cao, J., Zhang, Z., Luo, Y., Zhang, L., Zhang, J., Li, Z., & Tao, F. (2021). Wheat yield predictions at a county and field scale with deep learning, machine learning, and google earth engine. European Journal of Agronomy, 123, 126204. 10.1016/j.eja.2020.126204

Cheng, E., Zhang, B., Peng, D., Zhong, L., Yu, L., Liu, Y., Xiao, C., Li, C., Li, X., Chen, Y., Ye, H., Wang, H., Yu, R., Hu, J., & Yang, S. (2022). Wheat yield estimation using remote sensing data based on machine learning approaches. Frontiers in Plant Science, 13, 1090970. 10.3389/fpls.2022.1090970

Duguma, A.L., Bai, X. (2025). How the internet of things technology improves agricultural efficiency. Artificial Intelligence Review, 58, 63. 10.1007/s10462-024-11046-0

Fan, J., Li, Y., Yu, S., Gou, W., Guo, X., & Zhao, C. (2023). Application of Internet of Things to agriculture—The LQ-FieldPheno platform: A high-throughput platform for obtaining crop phenotypes in field. Research, 6, 0059. 10.34133/research.0059

Fan, J., Zhang, Y., Wen, W., Gu, S., Lu, X., & Guo, X. (2021). The future of internet of things in agriculture: Plant high-throughput phenotypic platform. Journal of Cleaner Production, 280, 123651. 10.1016/j.jclepro.2020.123651

Fan, Y., Liu, Y., Yue, J., Jin, X., Chen, R., Bian, M., Ma, Y., Yang, G., & Feng, H. (2024). Estimation of potato yield using a semi-mechanistic model developed by proximal remote sensing and environmental variables. Computers and Electronics in Agriculture, 223, 109117. 10.1016/j.compag.2024.109117

Farooq, M. S., Riaz, S., Abid, A., Umer, T., & Zikria, Y. B. (2020). Role of IoT technology in agriculture: A systematic literature review. Electronics, 9(2), 319. 10.3390/electronics9020319

Guo, F., Wang, P., Tansey, K., Zhang, Y., Li, M., Liu, J., & Zhang, S. (2024). A novel transformer-based neural network under model interpretability for improving wheat yield estimation using remotely sensed multi-variables. Computers and Electronics in Agriculture, 223, 109111. 10.1016/j.compag.2024.109111

Hochreiter, S., & Schmidhuber, J. (1997). Long short-term memory. Neural Computation, 9(8), 1735–1780. 10.1162/neco.1997.9.8.1735

Hoyos-Villegas, V., Jackson, M., Vargas-Cedeño, M., Farmer, E. E., Hanneman, M., Mazis, A., Singh, K. D., Sangjan, W., McNair, M., Sankaran, S., Tolosa, S. T., Gore, M. A., & Rife, T. W. (2025). Affordable phenomics: Expanding access to enhancing genetic gain in plant breeding. The Plant Phenome Journal, 8(1), e70034. 10.1002/ppj2.70034

Iserles, A. (1996). A first course in the numerical analysis of differential equations. Cambridge University Press.

Kasera, R.K., Gour, S., & Acharjee, T. (2024). A comprehensive survey on IoT and AI based applications in different pre-harvest, during-harvest and post-harvest activities of smart agriculture. Computers and Electronics in Agriculture, 216, 108522. 10.1016/j.compag.2023.108522

Liang, X., Yu, S., Ju, Y., Wang, Y., & Yin, D. (2025). Multi-scale remote-sensing phenomics integrated with multi-omics: Advances in crop drought–heat stress tolerance mechanisms and perspectives for climate-smart agriculture. Plants, 14(18), 2829. 10.3390/plants14182829

Liu, S., Ampatzidis, Y., Zhou, C., & Lee, W. S. (2025). AI-driven time series analysis for predicting strawberry weekly yields integrating fruit monitoring and weather data for optimized harvest planning. Computers and Electronics in Agriculture, 233, 110212. 10.1016/j.compag.2025.110212

Liu, S., Hu, Z., Han, J., Li, Y., & Zhou, T. (2022). Predicting grain yield and protein content of winter wheat at different growth stages by hyperspectral data integrated with growth monitor index. Computers and Electronics in Agriculture, 200, 107235. 10.1016/j.compag.2022.107235

Lu, J., Li, J., Fu, H., Zou, W., Kang, J., Yu, H., & Lin, X. (2025). Estimation of rice yield using multi-source remote sensing data combined with crop growth model and deep learning algorithm. Agricultural and Forest Meteorology, 370, 110600. 10.1016/j.agrformet.2025.110600

Marzougui, A., McGee, R. J., Van Vleet, S., & Sankaran, S. (2023). Remote sensing for field pea yield estimation: A study of multi-scale data fusion approaches in phenomics. Frontiers in Plant Science, 14, 1111575. 10.3389/fpls.2023.1111575

Numajiri, Y., Yoshino, K., Teramoto, S., Hayashi, A., Nishijima, R., Tanaka, T., Hayashi, T., Kawakatsu, T., Tanabata, T., & Uga, Y. (2021). IPOTs: Internet of Things-based pot system controlling optional treatment of soil water condition for plant phenotyping under drought stress. The Plant Journal, 107(5), 1569–1580. 10.1111/tpj.15400

Pathmudi, V. R., Khatri, N., Kumar, S., Abdul-Qawy, A. S. H., & Vyas, A. K. (2023). A systematic review of IoT technologies and their constituents for smart and sustainable agriculture applications. Scientific African, 19, e01577. 10.1016/j.sciaf.2023.e01577

Pineda-Castro, D., Diaz, H., Soto, J., & Urban, M. O. (2024). LysipheN: a gravimetric IoT device for near real-time high-frequency crop phenotyping: a case study on common beans. Plant Methods, 20, 39. 10.1186/s13007-024-01170-x

Pinto, F., Reynolds, M., & Schulthess, U. (2023). Satellite imagery for high-throughput phenotyping in breeding plots. Frontiers in Plant Science, 14, 1114670. 10.3389/fpls.2023.1114670

Rajak, P., Ganguly, A., Adhikary, S., & Bhattacharya, S. (2023). Internet of things and smart sensors in agriculture: Scopes and challenges. Journal of Agriculture and Food Research, 14, 100776. 10.1016/j.jafr.2023.100776

Ramírez-Arroyo, A., López, M., Rodríguez, I., Damsgaard, S. B., & Mogensen, P. (2025). Multi-connectivity solutions for rural areas: Integrating terrestrial 5G and satellite networks to support innovative IoT use cases. Smart Agricultural Technology, 12, 101260. 10.1016/j.atech.2025.101260

Sanaeifar, A., Guindo, M. L., Bakhshipour, A., Fazayeli, H., Li, X., & Yang, C. (2023). Advancing precision agriculture: The potential of deep learning for cereal plant head detection. Computers and Electronics in Agriculture, 209, 107875. 10.1016/j.compag.2023.107875

Sandhu, K. S., Mihalyov, P. D., Lewien, M. J., Pumphrey, M. O., & Carter, A. H. (2021). Combining genomic and phenomic information for predicting grain protein content and grain yield in spring wheat. Frontiers in Plant Science, 12, 613300. 10.3389/fpls.2021.613300

Sangjan, W., Carpenter-Boggs, L.A., Hudson, T. D., & Sankaran, S. (2022). Pasture productivity assessment under mob grazing and fertility management using satellite and UAS imagery. Drones, 6(9), 232. 10.3390/drones6090232

Sangjan, W., Carter, A. H., Pumphrey, M. O., Hagemeyer, K., Jitkov, V., & Sankaran, S. (2024). Effect of high-resolution satellite and UAV imagery plot pixel resolution in wheat crop yield prediction. International Journal of Remote Sensing, 45(5), 1678–1698. 10.1080/01431161.2024.2313997

Sangjan, W., Carter, A. H., Pumphrey, M. O., Jitkov, V., & Sankaran, S. (2021). Development of a Raspberry Pi-based sensor system for automated in-field monitoring to support crop breeding programs. Inventions, 6(2), 42. 10.3390/inventions6020042

Sangjan, W., Kick, D.R. & Washburn, J.D. (2025b) Improving plant breeding through AI-supported data integration. Theoretical and Applied Genetics, 138, 132. 10.1007/s00122-025-04910-2

Sangjan, W., McGee, R. J., & Sankaran, S. (2023). Evaluation of forage quality in a pea breeding program using a hyperspectral sensing system. Computers and Electronics in Agriculture, 212, 108052. 10.1016/j.compag.2023.108052

Sangjan, W., Pandey, P., Best, N. B., & Washburn, J. D. (2025a). MatchPlant: An open-source pipeline for UAV-based single-plant detection and data extraction. ArXiv. https://arxiv.org/abs/2506.12295

Segarra, J., Araus, J. L., & Kefauver, S. C. (2022). Farming and Earth observation: Sentinel-2 data to estimate within-field wheat grain yield. International Journal of Applied Earth Observation and Geoinformation, 107, 102697. 10.1016/j.jag.2022.102697

Shrestha, N., Powadi, A., Davis, J., Ayanlade, T. T., Liu, H., Tross, M. C., Mathivanan, R. K., Bares, J., Lopez-Corona, L., Turkus, J., Coffey, L., Jubery, T. Z., Ge, Y., Sarkar, S., Schnable, J. C., Ganapathysubramanian, B., & Schnable, P. S. (2025). Plot-level satellite imagery can substitute for UAVs in assessing maize phenotypes across multistate field trials. *Plants, People*, Planet, 7(4), 1011–1026. 10.1002/ppp3.10613

Silva, J. V., Heerwaarden, J. V., Reidsma, P., Laborte, A. G., Tesfaye, K., & Ittersum, M. K. V. (2023). Big data, small explanatory and predictive power: Lessons from random forest modeling of on-farm yield variability and implications for data-driven agronomy. Field Crops Research, 302, 109063. 10.1016/j.fcr.2023.109063

Sinha, B. B., & Dhanalakshmi, R. (2022). Recent advancements and challenges of Internet of Things in smart agriculture: A survey. Future Generation Computer Systems, 126, 169–184. 10.1016/j.future.2021.08.006

Skobalski, J., Sagan, V., Alifu, H., Al Akkad, O., Lopes, F. A., & Grignola, F. (2024). Bridging the gap between crop breeding and GeoAI: Soybean yield prediction from multispectral UAV images with transfer learning. ISPRS Journal of Photogrammetry and Remote Sensing, 210, 260–281. 10.1016/j.isprsjprs.2024.03.015

Tang, P., Liang, Q., Li, H., & Pang, Y. (2024). Application of internet-of-things wireless communication technology in agricultural irrigation management: A review. Sustainability, 16(9), 3575. 10.3390/su16093575

Tao, W., Zhao, L., Wang, G., & Liang, R. (2021). Review of the internet of things communication technologies in smart agriculture and challenges. Computers and Electronics in Agriculture, 189, 106352. 10.1016/j.compag.2021.106352

Tian, H., Wang, P., Tansey, K., Zhang, J., Zhang, S., & Li, H. (2021). An LSTM neural network for improving wheat yield estimates by integrating remote sensing data and meteorological data in the Guanzhong Plain, PR China. Agricultural and Forest Meteorology, 310, 108629. 10.1016/j.agrformet.2021.108629

Wang, H., Zhang, X., Liao, J., Zhang, Y., & Li, H. (2024). An improved adaptive data rate algorithm of LoRaWAN for agricultural mobile sensor nodes. Computers and Electronics in Agriculture, 219, 108773. 10.1016/j.compag.2024.108773

Xu, J., Gu, B., & Tian, G. (2022). Review of agricultural IoT technology. Artificial Intelligence in Agriculture, 6, 10–22. 10.1016/j.aiia.2022.01.001

Zhang, C., Marzougui, A., & Sankaran, S. (2020). High-resolution satellite imagery applications in crop phenotyping: An overview. Computers and Electronics in Agriculture, 175, 105584. 10.1016/j.compag.2020.105584

