## Supplementary Material S1 for "AGIcam: An Open-Source IoT-Based Camera System for Automated In-Field Phenotyping and Yield Prediction"

Summary of vegetation indices that were constructed in the study.

| Vegetation Index | Formulation | Reference |
|---|---|---|
| **CIgreen**: Chlorophyll Index Green | $\frac{NIR}{Green} - 1$ | Gitelson et al. (2003) |
| **EVI2**: Enhanced Vegetation Index 2 | $\frac{2.5 \times (NIR - Red)}{1 + NIR + (2.4 \times Red)}$ | Jiang et al. (2008) |
| **GNDVI**: Green Normalized Difference Vegetation Index | $\frac{NIR - Green}{NIR + Green}$ | Gitelson et al. (1998) |
| **NDVI**: Normalized Difference Vegetation Index | $\frac{NIR - Red}{NIR + Red}$ | Rouse et al. (1974) |
| **RDVI:** Renormalized Difference Vegetation Index | $\frac{NIR - Red}{\sqrt{(NIR + Red)}}$ | Roujean and Breon (1995) |
| **SAVI**: Soil Adjusted Vegetation Index | $\frac{1.5 \times (NIR - Red)}{NIR + Red + 0.5}$ | Huete (1988) |
| **SR:** Simple Ratio | $\frac{NIR}{Red}$ | Birth and McVey (1968) |

NIR: Near-Infrared; The ratio values of near infrared (NIR), red, green, and blue spectra of the Raspberry Pi's NoIR (no infrared) image from the Glenn & Tabb (2019) were used to transform the equation and used in this study:

CIgreen = $\dfrac{(0.666 \times Red)}{(0.233 \times Green)} - 1$

EVI2 = $\dfrac{(2.5 \times ((1.664 \times Blue) - (0.953 \times Red))}{(1 + (0.832 \times Blue) + (2.4 \times ((0.953 \times Red) - (0.832 \times Blue))))}$

GNDVI = $\dfrac{(1.664 \times Blue)}{(0.846 \times Green)} - 1$

NDVI = $\dfrac{(1.664 \times Blue)}{(0.953 \times Red)} - 1$

RDVI = $\dfrac{(1.664 \times Blue) - (0.953 \times Red)}{\sqrt{(0.953 \times Red)}}$

SAVI = $\dfrac{(1.5 \times \left((1.664 \times Blue) - (0.953 \times Red)\right))}{((0.953 \times Red) + 0.5)} - 1$

SR = $\dfrac{(0.832 \times Blue)}{((0.953 \times Red) - (0.832 \times Blue))}$

### References


Birth, G. S., & McVey, G. R. (1968). Measuring the color of growing turf with a reflectance spectrophotometer. *Agronomy Journal*, *60*(6), 640–643. https://doi.org/10.2134/agronj1968.00021962006000060016x

Gitelson, A. A., & Merzlyak, M. N. (1998). Remote sensing of chlorophyll concentration in higher plant leaves. *Advances in Space Research*, *22*(5), 689–692. https://doi.org/10.1016/S0273-1177(97)01133-2

Gitelson, A. A., Gritz †, Y., & Merzlyak, M. N. (2003). Relationships between leaf chlorophyll content and spectral reflectance and algorithms for non-destructive chlorophyll assessment in higher plant leaves. *Journal of Plant Physiology*, *160*(3), 271–282. https://doi.org/10.1078/0176-1617-00887

Glenn, D. M., & Tabb, A. (2018). Evaluation of five methods to measure normalized difference vegetation index (NDVI) in apple and citrus. *International Journal of Fruit Science*, *19*(2), 191–210. https://doi.org/10.1080/15538362.2018.1502720

Huete, A. (1988). A soil-adjusted vegetation index (SAVI). *Remote Sensing of Environment*, *25*(3), 295–309. https://doi.org/10.1016/0034-4257(88)90106-X

Jiang, Z., Huete, A. R., Didan, K., & Miura, T. (2008). Development of a two-band enhanced vegetation index without a blue band. *Remote Sensing of Environment*, *112*(10), 3833–845. https://doi.org/10.1016/j.rse.2008.06.006

Roujean, J., & Breon, F. (1995). Estimating PAR absorbed by vegetation from bidirectional reflectance measurements. R*emote Sensing of Environment*, *51*(3), 375–384. https://doi.org/10.1016/0034-4257(94)00114-3

Rouse, J. W., Jr., Haas, R. H., Schell, J. A., & Deering, D. W. (1973, December). *Monitoring vegetation systems in the Great Plains with ERTS* (Paper A-20, NASA SP-351, pp. 309–317). In Third Earth Resources Technology Satellite-1 Symposium, NASA Goddard Space Flight Center, Washington, DC.
