## Supplementary Material S2 for "AGIcam: An Open-Source IoT-Based Camera System for Automated In-Field Phenotyping and Yield Prediction"

### AGIcam Dashboard

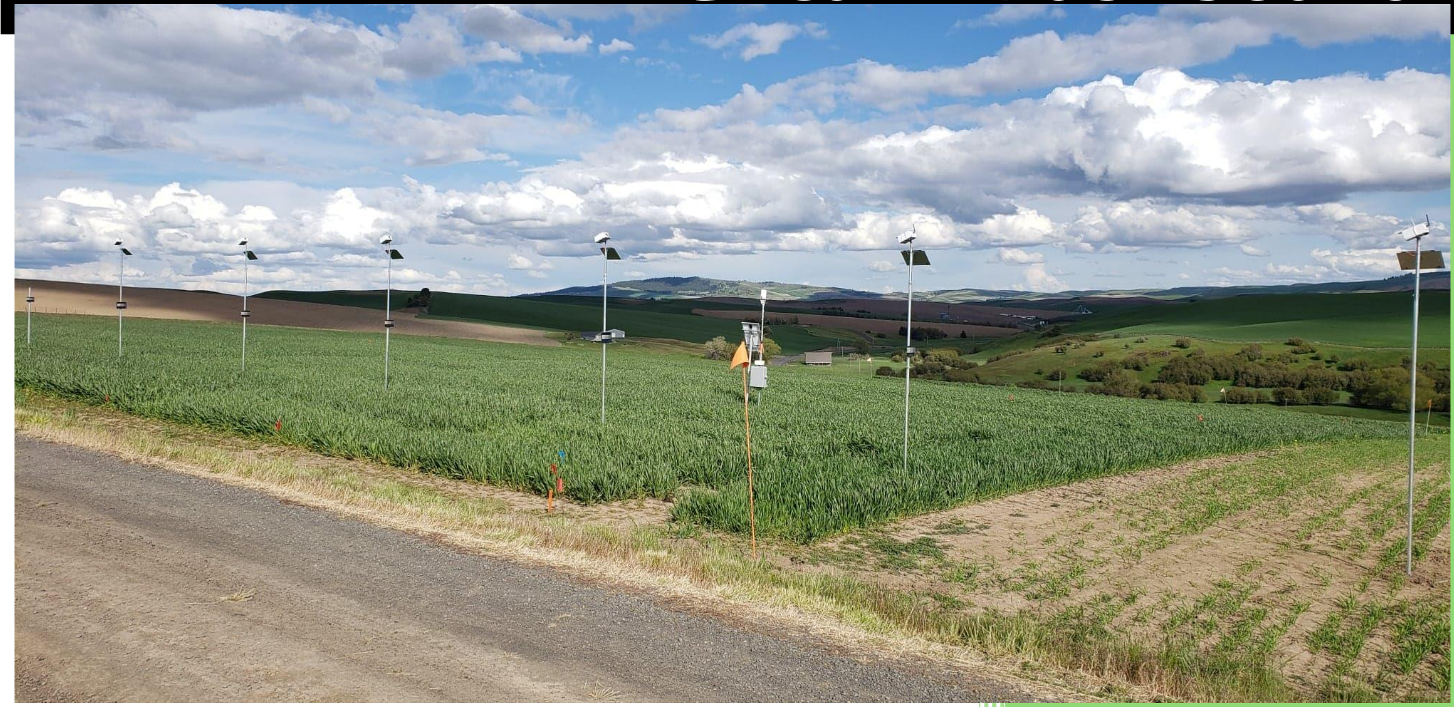

Worasit Sangjan

### Installation Map

The information and details of each AGIcam on the dashboard and the variety of trial plots, block number, etc., follow the map below (Figure 1):

- At the **Spring Wheat** field
    - 12 AGIcam were installed in the field
    - 3 plots were targeted with one AGIcam
- At the **Winter Wheat** field
    - 6 AGIcam were installed in the field
    - 4 plots were targeted with one AGIcam

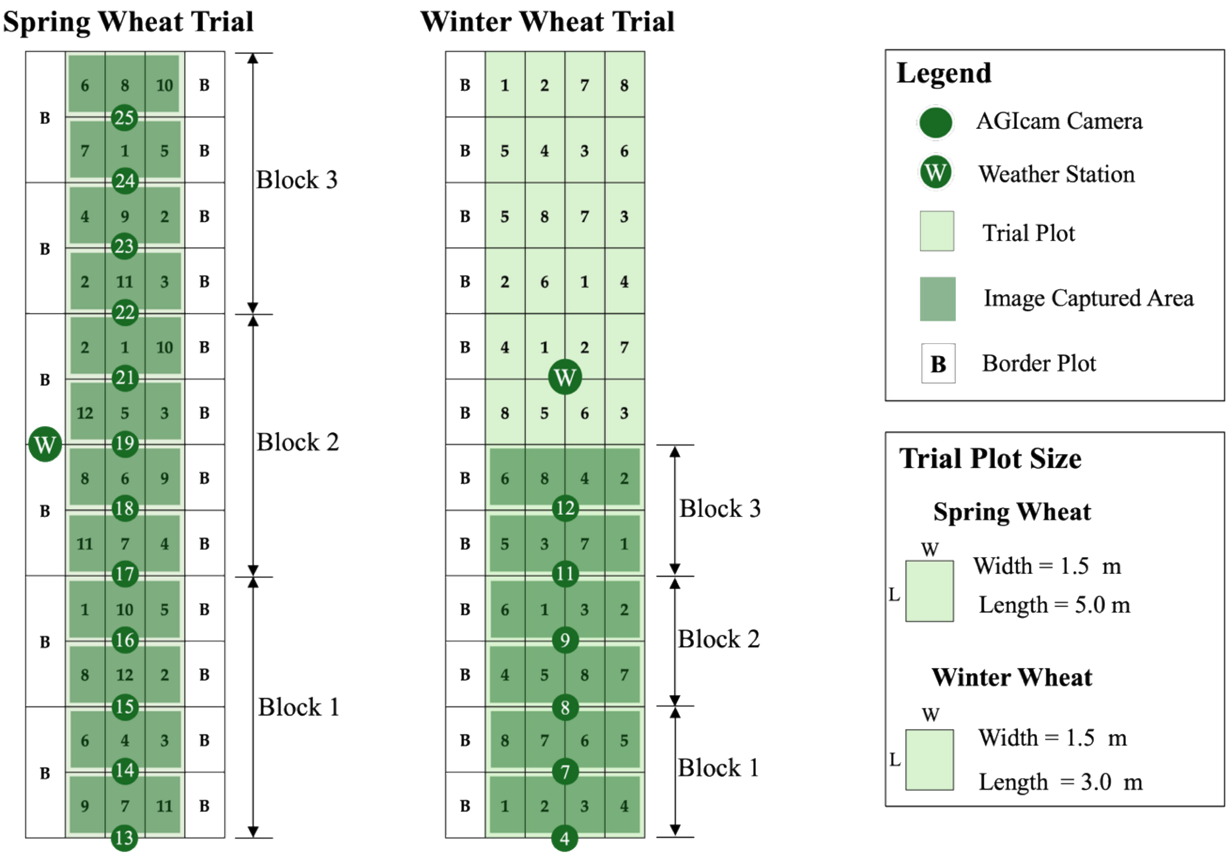


**Figure 1**. Plot layout and AGIcam system deployment for spring and winter wheat breeding trials during the 2022 season.

### AGIcam Dashboard

**URL**: The URL is currently unavailable, as the season has ended.

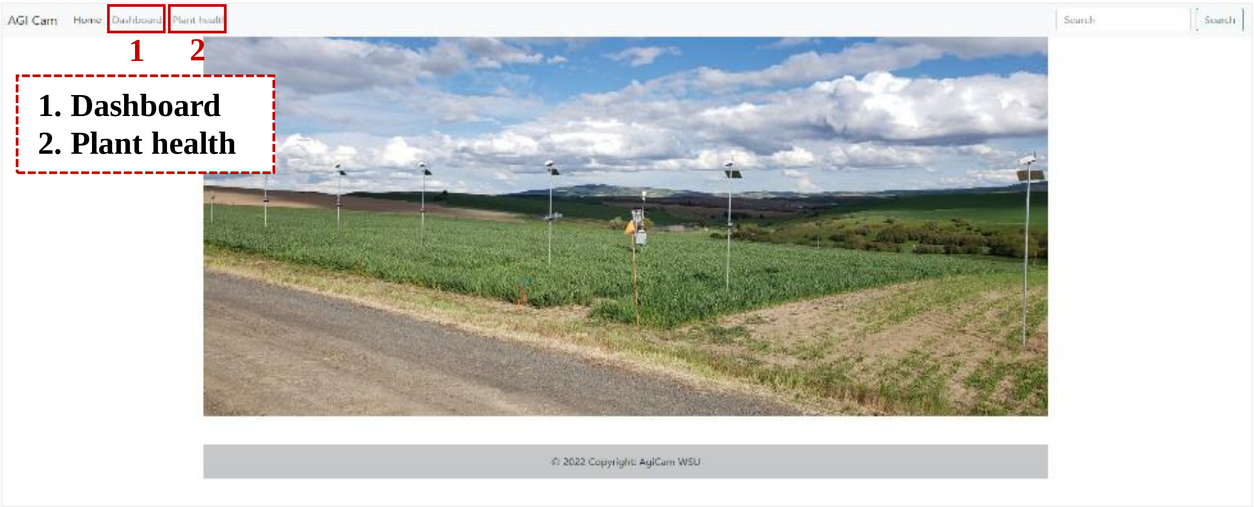


**Figure 2**. The AGIcam Dashboard's homepage. The source code for the dashboard is shared in GitHub (https://github.com/WorasitSangjan/IoT-based-Camera-Development).

1. **Dashboard** page: A list of all AGIcam to select to visualize the NDVI mean value.

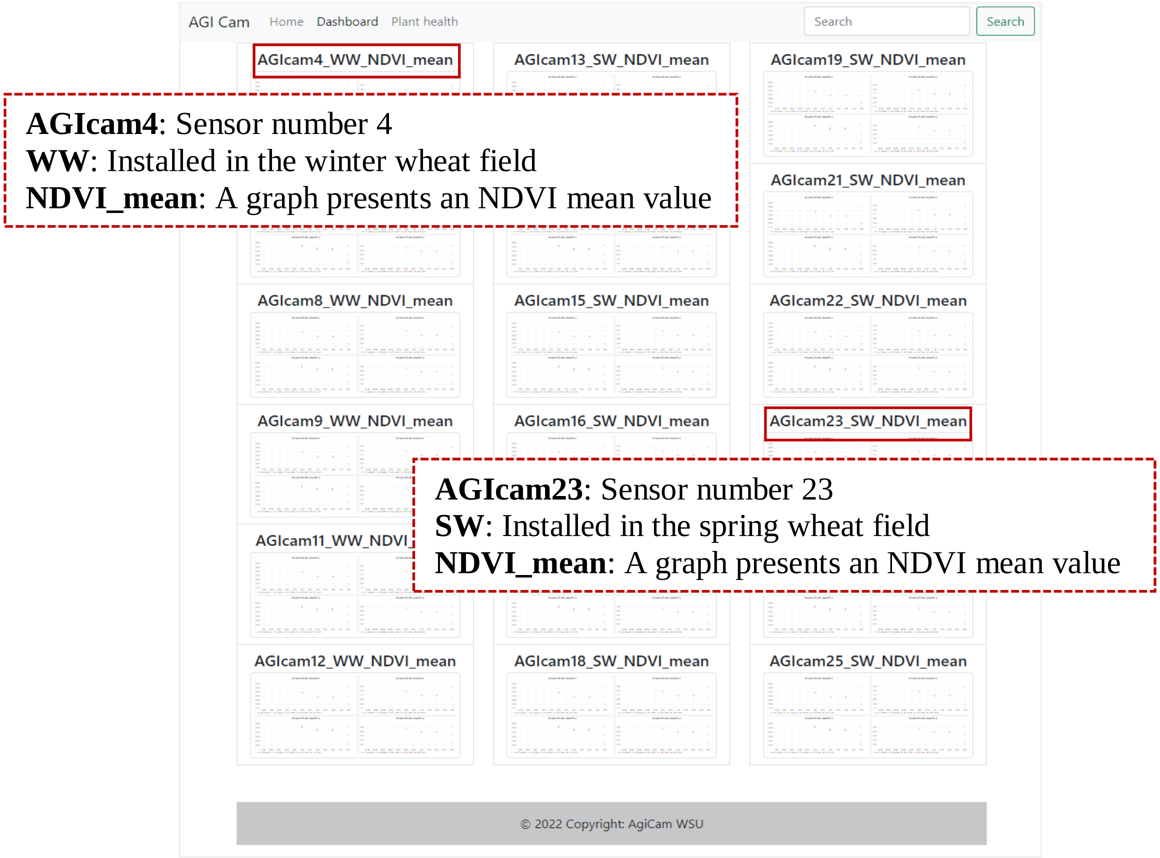


**Figure 3**. The Dashboard's page displays the list of AGIcam to check the NDVI value.

1.1 After clicking on each list item, the mean NDVI value of the corresponding individual plot will be displayed. Please refer to the map (Figure 1) to identify which AGIcam unit is associated with each plot.

- Figure 4 shows a mean NDVI of an individual plot after clicking “AGIcam4_WW_NDVI_mean.”
- The NDVI graphs of each trial plot are presented on this page. Four graphs are displayed, corresponding to AGIcam4, which monitors four plots of the 1st row in Block 1 (Varieties 1, 2, 3, and 4), as shown on the installation map (Figure 1).

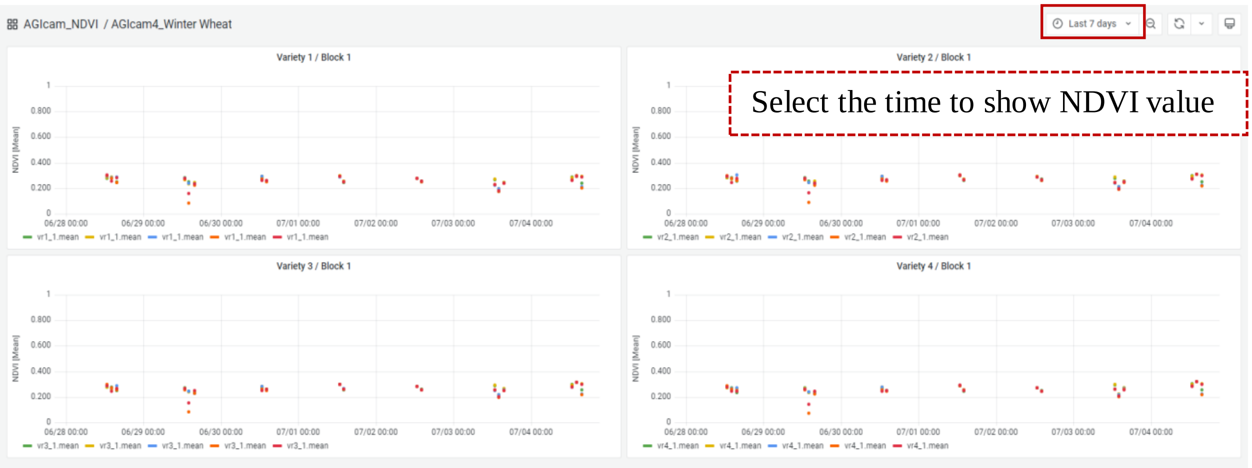


**Figure 4**. The mean NDVI graph of an individual plot.

1.2 The information inside each graph.

- Each AGIcam captures images three times a day (at 10:30 a.m., 12:00 p.m., and 1:30 p.m.), and at each time point, it captures five images.
    - Therefore, there are three periods of NDVI values per day, and at each period, there are five mean NDVI values.
- Each graph presents mean NDVI values for the last 7 days (default setting)
    - The time can be changed by clicking and selecting the tab in the right corner of the page (Figure 4).

2. **Plant health** page: A list of all AGIcam to select to visualize the captured images. Please refer to the map (Figure 1) to identify which AGIcam unit is associated with each plot.

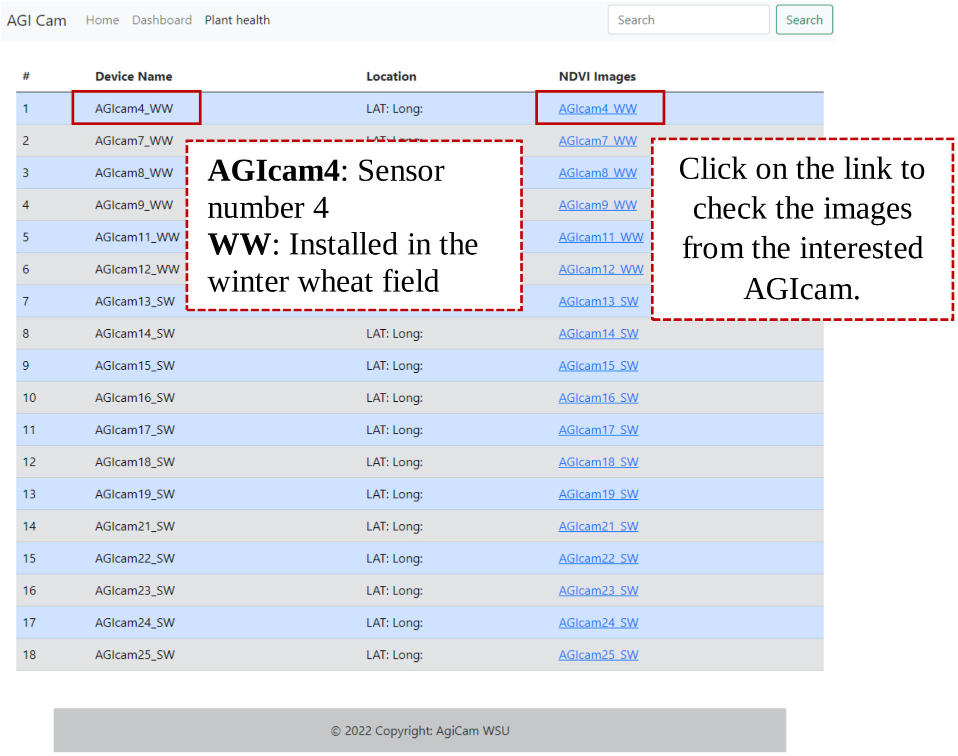


**Figure 5**. The Plant health's page displays a list of AGIcam to check the NDVI value.

2.1 After clicking on each list item, the captured image list for each day will be displayed.

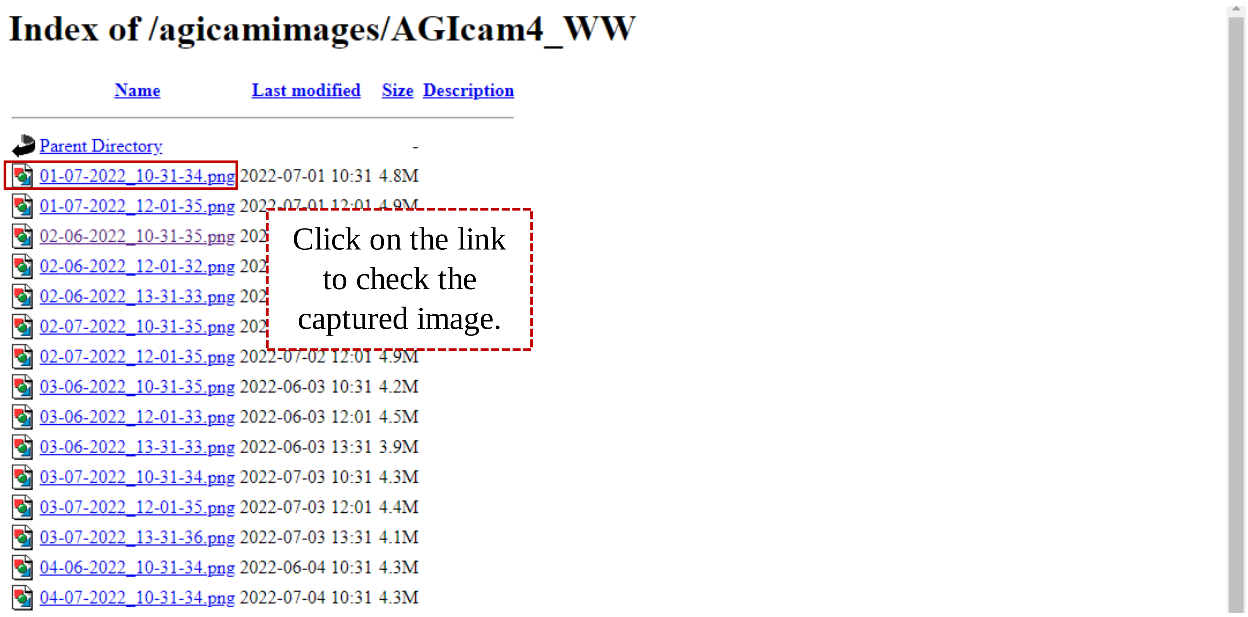

**Figure 6**. The page displays a list of images ordered by capture date and time.

2.2 The targeted trial plot.

- There are two images in one frame, an RGB image and an infrared image.
- The targeted plots of all AGIcam are the plots behind the reference panel.
- The image above is from AGIcam4; therefore, varieties 1, 2, 3, and 4 from the 1st row in Block 1 were captured (Figure 1)

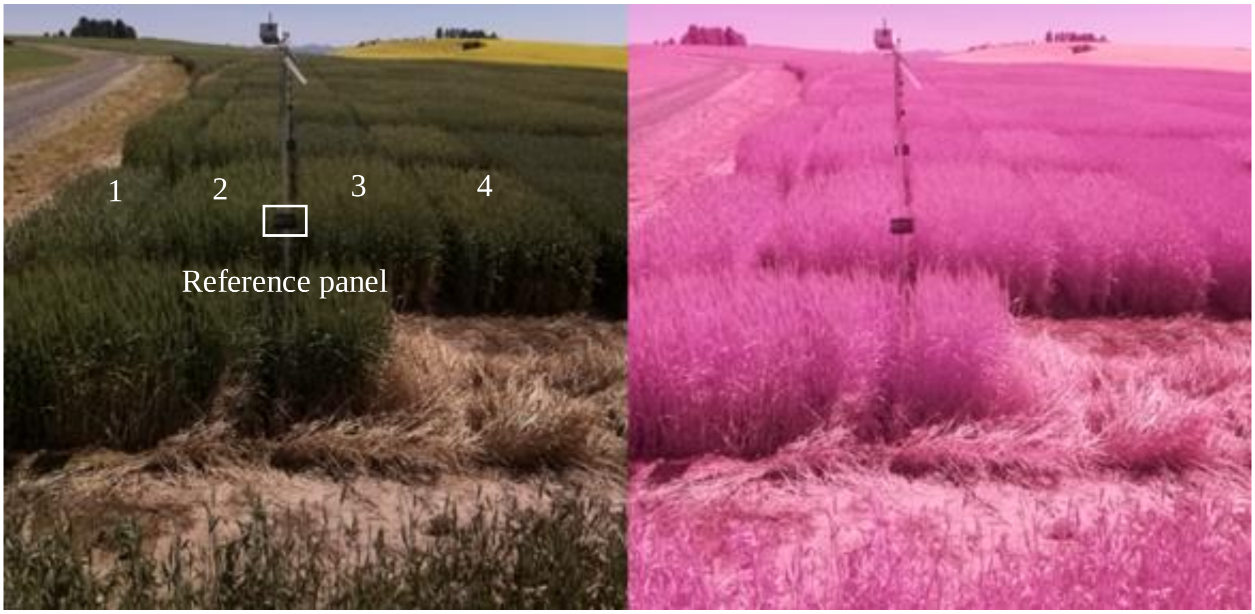


**Figure 7**. The captured image from AGIcam.

### AnyDesk

Worasit Sangjan

### AnyDesk: Remote Software

1. Install AnyDesk software on your computer (laptop, desktop) and/or smartphone.

- Download: https://anydesk.com/en
- Install the software on your computer.

2. Fill in the address of the remote targeted AGIcam

**Caution**:
- All AGIcams are set to capture images at 10:30 a.m., 12:00 p.m., and 1:30 p.m. Therefore, please do not remote access during these periods.
- Suggestion time for remote is 9:00–10:15 am and 3:00–4:30 pm, as the AGIcams are all turned on in these periods.

- After installing and opening the software, the page (Figure 1) will show.
- Fill in the address of the remote targeted AGIcam (see Table 1 for the address number).
- The software will ask for a Password to access the AGIcam: ton040298070

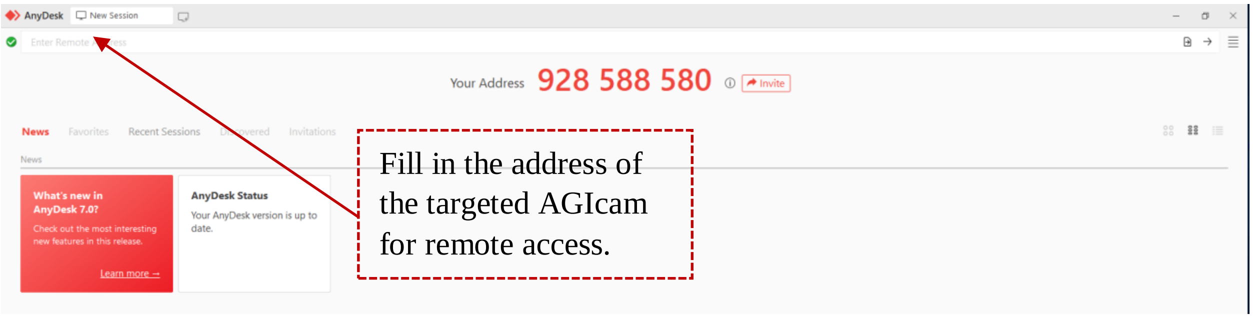


**Figure 1**. The AnyDesk homepage.

**Table 1**. Address numbers of the AnyDesk remote access software for each AGIcam.

| AGIcam | Address Number | AGIcam | Address Number |
|---|---|---|---|
| AGIcam4 | 654 062 980 | AGIcam16 | 311 678 612 |
| AGIcam7 | 249 651 876 | AGIcam17 | 206 685 061 |
| AGIcam8 | 899 347 374 | AGIcam18 | 415 249 812 |
| AGIcam9 | 176 395 825 | AGIcam19 | 896 234 517 |
| AGIcam11 | 243 131 173 | AGIcam21 | 725 132 815 |
| AGIcam12 | 333 564 996 | AGIcam22 | 616 505 174 |
| AGIcam13 | 976 708 099 | AGIcam23 | 813 809 273 |
| AGIcam14 | 247 920 114 | AGIcam24 | 859 983 777 |
| AGIcam15 | 844 623 462 | AGIcam25 | 271 416 515 |

3. Access to check the images or the algorithms on AGIcam.

- A storage drive is available on the desktop of all AGIcam (Figure 2). The user can navigate to the storage and check the images in the folder named "image."
- Algorithms are in the drive C.

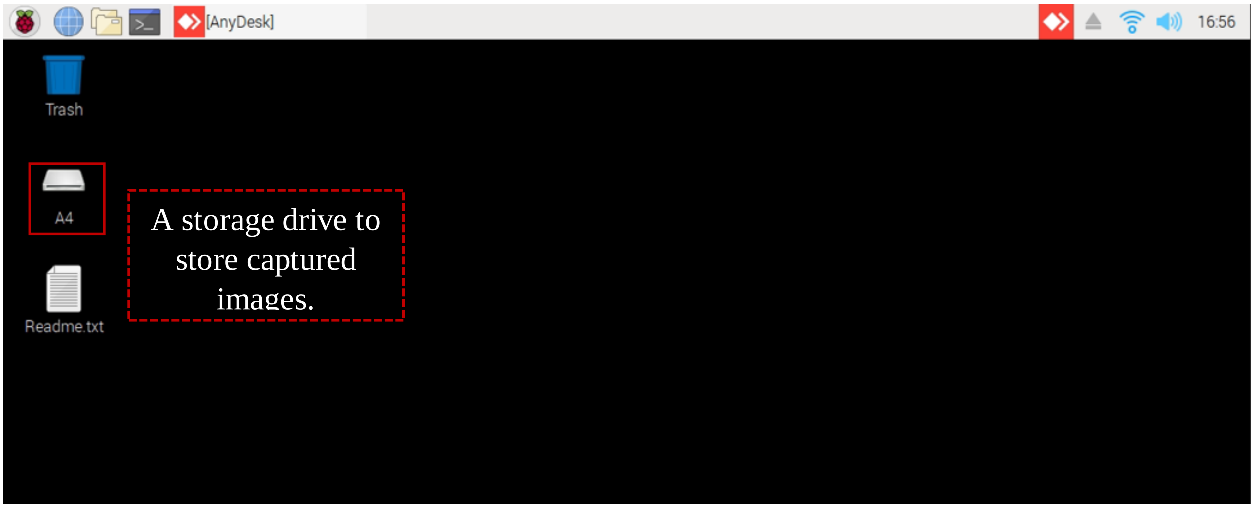


**Figure 2**. Desktop interface of the AGIcam (Raspberry Pi)
